# A mixture of plant polyphenols unexpectedly aggravates liver metastasis of colorectal cancer in mice

**DOI:** 10.64898/2026.02.15.703962

**Authors:** Merve Erdem, Johanna Roth, Jana Knobloch, Jochen Nolting, Hannes Hatten, Eray Sahin, Felix Schön, Sandra Halbfeld, Nicole S. Treichel, Thomas Clavel, Roman Bülow, Christian Liedtke, Thorsten Cramer

**Author notes:** Corresponding author: Thorsten Cramer, Molecular Tumor Biology, Department of General, Visceral, Pediatric and Transplantation Surgery, Uniklinik RWTH Aachen, Pauwelsstraße 30, 52074 Aachen, Germany. Department of Pediatrics, University Hospital Würzburg, Würzburg, Germany (Eray Sahin).

## Abstract

Epidemiological studies suggest that vegetarian diets are associated with lower cancer incidence and mortality, an effect attributed in part to phytochemicals such as polyphenols and carotenoids. Although numerous in vitro experiments and investigations using immunodeficient rodent models report tumor-suppressive activities of phytochemicals, their impact on tumor progression in immunocompetent hosts remains insufficiently understood. Here, we examined the influence of a defined plant phytochemical mixture (PPM) on the growth of colon cancer liver metastases, both *in vitro* and in immunocompetent mice. Consistent with the prevailing literature, treatment of the murine colon cancer cell line MC38 with the PPM significantly reduced cell proliferation and survival *in vitro*. Strikingly, however, administration of the PPM to mice bearing MC38-derived hepatic metastases markedly accelerated tumor growth. Immunohistochemical analyses revealed a significantly increased accumulation of immune cells—specifically CD45⁺ leukocytes and F4/80⁺ macrophages—at the periphery of the metastatic lesions in PPM-treated animals. To assess the functional relevance of this inflammatory response, the PPM was combined with the anti-inflammatory drug prednisolone. This intervention resulted in significantly reduced metastatic burden, supporting the notion that the PPM exacerbates tumor progression through enhanced peritumoral inflammation. These findings highlight the importance of validating observations from cell culture and immunodeficient models in fully immunocompetent systems. They further emphasize that the immunomodulatory effects of plant phytochemicals warrant careful and comprehensive investigation.

## Introduction

Colorectal cancer (CRC) ranks as the third most frequent malignant tumor worldwide and is accompanied by substantial cancer-associated morbidity and mortality [1]. While the prognosis of patients with locally confined tumor growth has improved significantly in recent years [2], clinical care of metastasized CRC remains challenging [3]. Due to continuous improvement of surgical techniques and establishment of interdisciplinary therapy planning, CRC patients with liver metastases can nowadays undergo treatment with curative intent [4]. This is an impressive achievement and about 25% of operated patients survive five years or longer [4]. Unfortunately, the majority of patients still develop tumor recurrence in the liver or other organs and will, ultimately, succumb to the disease [2]. This demonstrates the urgent need to identify innovative approaches to target colorectal cancer liver metastasis.

Diets that are characterized by the exclusion or reduction of meat and other animal products, e.g. vegetarian and vegan diets, have gained significant attention for their potential health benefits and role in disease prevention. Epidemiological studies have increasingly investigated the incidence and risk of various cancers among vegetarians compared to omnivores, often suggesting a lower overall cancer risk in those adhering to plant-based diets [5, 6]. It is largely hypothesized that this protective effect is achieved by higher intakes of plants rich in bioactive compounds such as fruits, vegetables, whole grains, and legumes [7]. Phytochemicals—bioactive compounds found in plants such as polyphenols, flavonoids, and carotenoids— exert potent antioxidant, anti-inflammatory, and anti-proliferative properties, contributing to the modulation of cellular processes involved in carcinogenesis [8]. This led to the assumption that a better understanding of the interplay between vegetarian dietary patterns, epidemiological cancer trends, and the mechanistic roles of plant polyphenols will offer valuable insights into strategies for both cancer prevention and therapy.

In rodent model systems the inhibitory effect of phytochemicals on the trajectory of cancer is well established. Mechanistically, these compounds have been shown to reduce DNA damage caused by carcinogens, enhance the activity of detoxifying enzymes, and modulate signaling pathways involved in cell proliferation, apoptosis, and inflammation [7, 9]. For example, flavonoids like quercetin and catechins have suppressed tumor formation in chemically induced cancer models by interfering with mutagenesis and promoting cancer cell death [10, 11]. Similarly, curcumin and resveratrol have been found to inhibit tumor growth and metastasis by targeting multiple molecular mechanisms, including inhibition of angiogenesis [12] and modulation of immune responses [13]. Clinical translation of these findings is challenged by the fact that the concentrations at which most phytochemicals display the greatest efficiency *in vitro* are too high to be achieved *in vivo*. This is most importantly explained by the low bioavailability of phytochemicals after dietary consumption or oral gavage of the isolated substances [14]. It was reasoned before that combining different polyphenols compared to application of the individual compounds would be more effective *in vivo* [15, 16]. It is well accepted in oncology that combined targeting of different pro-tumorigenic pathways enhances the overall anti-tumorigenic response [17]. This approach was further supported by published work from the laboratory of Stephen Hursting showing that oral consumption of a mixture of 12 different fruits and vegetables significantly inhibited intestinal tumorigenesis in Apc^Min^ mice [18]. Against this background, we decided to test the efficacy of a plant polyphenol mixture in a well-established model of colon cancer liver metastasis in immunocompetent mice [19].

## Materials and Methods

### Preparation of the plant polyphenol mixture (PPM)

The ingredients of the PPM were carefully selected based on published results [20] from Richard Béliveau and colleagues (Université du Québec à Montréal, Montréal, Canada), international pioneers regarding the role of phytochemicals for tumor prevention. The composition of the PPM is shown in table S1. Vegetables were purchased from a local organic food market, thoroughly cleaned with tap water, measured, and added to a juicer (Philips Spin Juicer, HR1855/70) one at a time. Then, the long pepper was added to the mixture and blended. The garlic was peeled, crushed, and added to the mixture. Turmeric powder was first mixed with linseed oil and later added to the mixture. Finally, the green tea polyphenols were added and thoroughly mixed. The PPM was centrifuged at 50 000 x *g* at 4°C for 45 minutes. The supernatant was collected, aliquoted, and stored at −80 °C. Sterile filtration (0,22 µm) was performed immediately before using the PPM *in vitro* and *in vivo*.

### Cell culture

MC38 murine colon carcinoma cells [21] were kindly provided by Dr. Raffaella Giavazzi (Istituto di Ricerche Farmacologiche Mario Negri, Milano, Italy) and grown in Dulbecco’s Modified Eagle’s Medium Ham’s F-12 Nutrient Mixture (Sigma-Aldrich, D8437) supplemented with 10% fetal bovine serum (Thermo Fisher Scientific, A5256701) and 100 U/ml penicillin, 100 μg/ml streptomycin (Sigma-Aldrich, P0781) in a humidified incubator with 5% CO_2_ at 37°C. Cells at passages 8-15 were used for the experiments. MC38 cells were authenticated using highly polymorphic short tandem repeat loci (STRs) [22] by Multiplexion (Friedrichshafen, Germany).

### Cell proliferation assay

For cell proliferation experiments, MC38 cells were seeded in 6-well cell culture plates at 1.5 × 10^5^ cells/well. On the next day, different dilutions of the PPM were prepared with cell culture media and applied to the cells (2 ml/well). After 48 hours, cell images were taken using a Leica DM IL LED microscope with DFC345 FX at 10x magnification, followed by cell trypsinization (Pan Biotech, P10-023100) and cell counting (LUNA-FL™ Dual Fluorescence Cell Counter).

### Flow cytometry-based analysis of cell proliferation and cell cycle distribution

The effect of the PPM on cell proliferation was investigated using two flow cytometric procedures. The first technique measures the fluorescence of intracellular amines, covalently stained prior to PPM addition, whose concentration and intensity per cell decreases with each cell division. With the second method the cell cycle state was determined by DNA staining with propidium iodide. The flow cytometer used was an LSR Fortessa (Becton Dickinson). Intracellular amines were stained using the commercial *CellTrace Cell Proliferation Kit* (Invitrogen #C34570), containing the fluorescent dye CFSE (carboxyfluorescein succinimidyl ester), according to the manufacturer’s instructions. Briefly, 1 × 10^5^ cells per well of a 6-well cell culture plate were incubated for one day in 4 ml of cell culture medium. After removing the medium, 1 ml of PBS supplemented with 5 µM CFSE was added. Following 20 minutes incubation, the PBS-CFSE solution was replaced with 4 ml of fresh cell culture medium. PPM was added at a dilution of 1:100. Control wells remained untreated. Control samples were measured on day 0 and day 3, PPM samples were measured on day 3 only. For flow cytometry, the medium was removed, the wells were then washed with PBS, trypsinized, washed again, and finally the cells were collected in PBS. Fluorescence excitation was performed at 488 nm, detection at 515 ± 15 nm. DNA staining for cell cycle analysis was performed using a protocol established in our laboratory with the ready-to-use *Propidium Iodide (PI)/RNase Staining Solution*, Cell Signaling Technology, 4087). 1.5 × 10^5^ cells per well of a 6-well plate were incubated overnight in 4 ml of medium. The next day, the medium was replaced with a medium containing PPM at ratios of 1:250, 1:175 or 1:100 or with cell culture medium only (controls). After two days, the medium was removed, cells were trypsinized, washed and resuspended in 1 ml of PBS. 100 µl of this cell suspension was taken for cell counting in the Luna Cell Counter, and the remaining 900 microliters were used for cell cycle analysis. To stain with PI, the cells were fixed with cold ethanol at −20°C for at least 20 minutes. After centrifugation at 104 x *g* for 5 minutes, the supernatant was discarded. The cell pellet was washed in PBS and PI staining was then performed by adding 300 µL of PI/RNase staining solution. After incubating at 37°C for at least 20 minutes, flow cytometric measurements were taken with an excitation wavelength of 561 nm and fluorescence detection at 610 ± 10 nm.

### Generation of MC38-FUCCI cells

In order to monitor and visualize cell cycle progression in MC38 cells in higher timely resolution by time lapse microscopy, we integrated an established cell cycle reporter construct referred to as FUCCI [23] into the ROSA26 locus using CRISPR-Cas9-based methodology. To this end, a donor plasmid was designed containing two ROSA26-specific homology arms of approximately 800 bp each as well as a bicistronic FUCCI cassette under control of the EF-1α promotor. The donor plasmid was generated by VectorBuilder (Neu-Isenburg, Germany). Donor and gRNA were transfected into MC38 cells using Lipofectamine 3000 (Thermo Fisher Scientific) and standard procedures. Transfected single cell clones were isolated using a BD FACSAria™ Fusion 5 Flow Cytometer (Becton Dickinson, New Jersey, USA) and checked for targeted integration of the FUCCI reporter by locus-specific PCR. Further details on the donor plasmid, ROSA26-specific gRNA and primers will be made available upon reasonable request.

### Analysis of cell cycle dynamics and nuclear size in MC38-FUCCI cells

MC38-FUCCI cells were seeded into 6-well culture plates at a density of 5 × 10^4^ per well and left untreated for 24 hours. The medium was replaced with a medium containing PPM at ratios of 1:100, 1:200 or full medium without PPM serving as control. Subsequently, the plates were transferred into a Axio Observer.Z1/7 microscope with an incubation unit (Carl Zeiss AG, Oberkochen, Germany). Incubation during the live cell imaging over the course of 48 hours was done at 37°C and 5% CO_2_. For each condition, 16 images for every channel were taken every 15 minutes. Each image stack was imported into the open source Bioimage Analysis software QuPath [24] and individually analyzed for cell count, nucleus size and cell cycle phase. Statistical analysis was run in the integrated development environment RStudio [25].

### Generation and imaging of MC38-FUCCI derived spheroids

4 × 10^3^ MC38-FUCCI cells per well were seeded into super low attachment 96-well culture plates (Sarstedt, Nümbrecht, Germany) and grown for 48 hours. Successful formation of MC38-FUCCI spheroids was confirmed via brightfield microscopy. The medium was replaced with a medium containing PPM at ratios of 1:100, 1:200 or no PPM (negative control). Subsequently, the plates were transferred into a Axio Observer.Z1/7 microscope with incubation unit (Carl Zeiss AG, Oberkochen, Germany). Incubation during the live cell imaging was done at 37°C and 5% CO_2_ for up to 72 hours. One image for every channel was taken every 15 minutes.

### Animal experiments

All animal experiments were approved by the governmental committee, LANUK (Landesamt für Natur, Umwelt und Klima, Nordrhein-Westfalen, Recklinghausen, Germany) with the reference number of 81-02.04.2017.A398 and performed in accordance with federal German law and the *Guide for the Care and Use of Laboratory Animals* (8th edition, NIH publication, 2011, USA). C57BL/6J mice (8-12 weeks old) were used in the experiments. The animals were housed in a specific pathogen-free animal facility and maintained with *ad libitum* access to water and standard rodent chow (Rat/Mouse Maintenance, ssniff, V1534-300) in a gastight room under strict 12h light/dark cycles (day: 7:00 am-7:00 pm) at 22 ± 1°C room temperature.

### Murine model of colorectal cancer (CRC) liver metastasis

Each surgical procedure was performed under general anaesthesia. Analgesia was administered by subcutaneous injection of buprenorphine (0.05 mg/kg BW) 30 minutes before surgery and at 8-hour intervals for 3 days postoperatively. During surgery, anesthesia was maintained with an isoflurane-oxygen mixture (induction: 5% isoflurane, continuation: 1-1.5% isoflurane, oxygen flow 2l/min), and mice were placed on a heating pad in a horizontal position. Through a midline laparotomy in the abdominal cavity, a retractor system was placed on the peritoneum. Tumor cell suspension (7.5 × 10^4^ MC38 cells in 250 µl Hank’s balance salt solution (Thermo Fisher Scientific, #14175053)) was injected into the spleen in 15 seconds using a 29G cannula, and the needle was left in the tissue for an additional 2 minutes to prevent bleeding. The hilar vessels were ligated using electrocautery and the spleen was immediately removed (splenectomy). The abdominal wall was closed by suturing first the peritoneum, followed by the skin layer. 14 days after surgery, the experimental group was administered with 100 µl PPM daily via oral gavage for 7 days. All mice were sacrificed 21 days after operation and liver tissues, blood serum and cecum stool samples were collected (Fig. S1A). Tissue samples were snap frozen and liver tissues were fixed in 10% formalin overnight, followed by paraffin embedding next day. Blood samples were collected both into sample tube EDTA K3E (Sarstedt, Germany) for blood count analysis and into serum-gel Z tubes (Sarstedt, Germany) for liver damage marker analysis. Serum-gel Z tubes were allowed to clot for 30 minutes at room temperature and centrifuged at 10 000 x *g* for 5 minutes. Serum and blood samples were directly given to the Biochemistry Laboratory of Institute for Laboratory Animal Science, University Hospital RWTH Aachen. To inhibit the inflammatory response, mice received intraperitoneal (i.p.) injections of prednisolone (Prednisolut® 25 mg L, MIBE GmbH, Sandersdorf-Brehna, Germany) at a final concentration of 10 mg/kg of body weight [26] in addition to the PPM gavage (Fig. S1B). For the experiments with metastasis-free (healthy) mice, animals were treated with 100 µl PPM daily via oral gavage for 7 days and then sacrificed (Fig. S1C). Blood and tissue samples were processed as described above.

### Isolation of total RNA and quantitative real-time PCR analysis

Total RNA was extracted from frozen tumor tissues using TRIzol reagent (Invitrogen, 15596026) according to the manufacturer’s instructions. RNA quality and concentration were determined via UV spectrophotometry in Synergy HT Multi-Mode Microplate Reader (BioTek, USA). RNA yields ranged between 600-1200 ng and A_260_/A_280_ ratios were between 1.8-2. Genomic DNA was removed using RNase-free DNase I treatment (Thermo Fisher Scientific, AM2222), and 1 µg total RNA of each sample was reverse transcribed using iScript cDNA Synthesis Kit (Bio-Rad, 1708891) in line with the kit instructions. Quantitative real-time PCR analysis was performed on ABI 7500 Real-Time PCR system using *Power* SYBR Green PCR Master Mix (Applied Biosystems, 4367660). The primer sequences are provided in the Supplementary Methods section. Primer-specific annealing temperatures were optimized beforehand, and product specificity was checked by melt curve analyses followed by agarose gel electrophoresis. Amplification efficiency was calculated in LinRegPCR using raw data (Heart Failure Research Center, Amsterdam, The Netherlands) [27], and relative mRNA expression levels were calculated with the comparative delta-CT method and normalized to the housekeeping gene *B2m* [28].

### Immunohistochemistry

All stainings were performed on formalin-fixed, paraffin-embedded tissues beginning with deparaffinization followed by rehydration of the sections. For hematoxylin and eosin (H&E) staining, the slides were first stained in hematoxylin solution for 5 minutes, washed with warm tap water, incubated with 1% eosin for 5 seconds and subsequently rinsed in acetic acid solution. For immunohistochemistry (IHC), antigen retrieval was done by 15 minutes of heating at 110°C in target retrieval solution (Dako, S1699) using the Decloaking Chamber (Biocare Medical, USA). The slides were washed twice with TBST (Dako, S3306), incubated with 3% H_2_O_2_-PBS solution for 10 minutes to inactivate endogenous peroxidase and blocked with 2.5% normal goat serum (ImmPress HRP anti-rat IgG Kit, Vector Laboratories, MP-7444-15) for 10 minutes. Slides were then incubated overnight at 4°C with either CD45 (Thermo Fisher Scientific, #14-0451-82) or F4/80 (Thermo Fisher Scientific, #14-4801-82) antibodies, both diluted 1:1.000 in “Dako Antibody Diluent” (Dako, S3022). Next day, secondary antibody incubation (ImmPress HRP anti-rat IgG Kit, Vector Laboratories, MP-7444-15) for 30 minutes at room temperature and application of DAB solution (Zytomed, DAB-057) was performed. The reaction was stopped in distilled water, and counterstaining of the nuclei was done with hematoxylin. Representative images of IHC stainings were captured using the Zeiss Axio Imager.M2 microscope equipped with the Axiocam 503 Color camera.

### Quantification of immunohistochemistry images

Scanning of the IHC images were carried out using the Vectra 3 Automated Quantitative Pathology Imaging System (Akoya Biosciences). Segmentation and counting analyses of the whole image areas were carried out using QuPath version 0.4.3 [24]. Briefly, after the annotation for liver and tumor areas, a cell detection algorithm was used to detect cell structures based on nuclei, and later the pixel classifier tool was run to precisely detect positively stained areas. After determining the number of total cells and positive objects, the percentage of positive cells was calculated for each sample.

### 16S rRNA gene amplicon sequencing and analysis

Metagenomic DNA extraction from gut samples, library preparation and sequencing were performed by the Functional Microbiome Research Group at Uniklinik RWTH Aachen. Briefly, frozen cecal stool samples were used to isolate metagenomic DNA, following a modified protocol from Godon *et al.* [29]. The V3-V4 regions of 16S rRNA genes were amplified in duplicate, using primers 341F-785R [30] in a two-step PCR protocol (15 + 10 cycles) [31] implemented within an automation platform (Biomek400, Beckman Coulter). After purification with AMPure XP system, amplicon libraries were sequenced using an Illumina MiSeq system in paired end mode (2×300nt) using single barcodes according to the manufacturer’s instructions. Raw reads were quality filtered using fastp [32] and imported into Qiime2 [33] environment to produce amplicon sequence variants (ASVs) using DADA2 workflow [34]. ASVs were taxonomically annotated using mouse stool weighted SILVA classifier [35]. Rooted phylogenetic tree, ASV relative abundance table, and taxonomic information of the ASVs were exported and used in downstream analyses [36]. Alpha and beta diversity, differential abundance (DA), and statistical analyses with respective visualizations were carried out in R. Detailed analysis steps were described in the Supplementary Methods.

### Statistical analyses

Graphic display and statistical analyses were conducted using GraphPad Prism (Version 10.6, California, USA). Illustrations in figure 7 and figure S1 were created using BioRender (https://biorender.com). Data are presented as mean with standard deviation (SD) unless otherwise indicated. The data were first subjected to a normalization test, and significance values were calculated using Student’s unpaired t-test or Mann Whitney U test when comparing two groups. For other analysis, one-way or two-way analysis of variance (ANOVA) with appropriate post-hoc test was performed. *p* values less than 0.05 were considered statistically significant and are represented as * *p* < 0.05, ** *p* ≤ 0.01, *** *p* ≤ 0.001, **** *p* ≤ 0.0001.

### Data Availability

The raw sequencing data files used in microbiota analysis were submitted to the European Nucleotide Archive and are publicly available with the accession number PRJEB108032.

## Results

### The plant polyphenol mixture (PPM) dose-dependently inhibits proliferation of murine colon cancer cells *in vitro*

We initiated our analysis by incubating the well-established and widely used murine colon cancer cell line MC38 [21] with different concentrations of the PPM. As can be seen in figure 1A and B, the PPM dose-dependently inhibited the survival of MC38 cells in cell culture. Next, cell proliferation was investigated more specifically using CFSE (carboxyfluorescein succinimidyl ester, for details see Materials and Methods). Figure 1C demonstrates a broad appearance of CFSE-positive cells after addition of the PPM compared to control, supporting an inhibitory effect of the PPM on cell proliferation. To further characterize the effect of the PPM, cell cycle distribution was analysed via flow cytometry after incubation with propidium iodide. This demonstrated multiple effects of the PPM, most prominently increased <2n, reduced 2n and increased >4n fractions (Fig. 1D and S2A). Cells in >4n typically have increased DNA content, multiple chromosomal aberrations and have been termed “polyploid giant cancer cells” (PGCC) [37]. Treatment of MC38 with the PPM resulted in the appearance of large cells with multiple nuclei (not shown), morphologically resembling PGCCs and therefore nicely complementing the FACS data.

**Figure 1.**
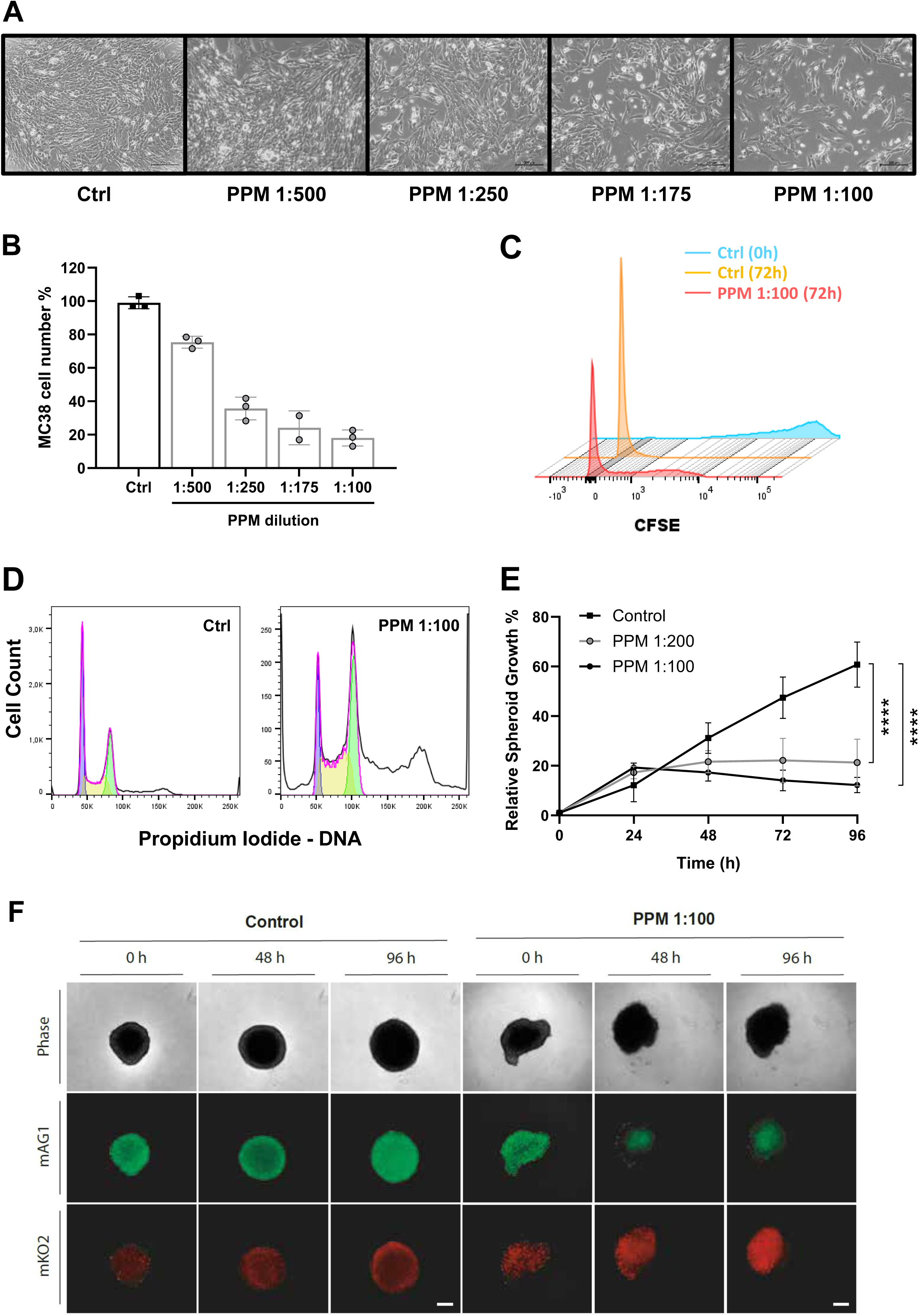
Effect of the plant polyphenol mixture (PPM) on MC38 cells *in vitro*. **A.** Representative images of Ctrl and PPM-treated MC38 cells at the 48h time point (scale bar: 200 µm). **B.** Proliferation of MC38 cells treated with the PPM at 1:500, 1:250, 1:175, and 1:100 dilutions for 48h. Cell numbers were normalized to the control (Ctrl) samples and presented as percentages. **C.** Flow cytometry results of MC38 cells stained with CFSE dye to trace cellular proliferation. The yellow peak is Ctrl and red peak is PPM (1:100 dilution)-treated MC38 cells at 72h. The blue peak represents MC38 Ctrl cells immediately after CFSE staining. The y-axis (not shown) represents the cell counts. **D.** Cell cycle analysis of MC38 cells using flow cytometry and propidium iodide staining. Plots are shown for Ctrl and PPM (1:100 dilution)-treated MC38 cells after 48h. Purple, yellow, and green areas correspond to G1, S, and G2/M phases, respectively. **E.** Spheroids generated from MC38-FUCCI cells were seeded in super-low-attachment plates and treated with PPM at the indicated concentrations. Relative spheroid growth in diameter was measured every 24 hours over the course of 96 hours. Relative numbers were normalized to baseline measurements of each treatment condition at the beginning of the experiment (0 hours). Data are displayed as mean ± SD. Statistical significance was determined using Dunnett’s multiple comparisons test, comparing PPM treatment to control (\*\*\**p* ≤ 0.0001). Image quantification was achieved from *n* = 8 spheroids. **F.** Representative images of MC38-FUCCI spheroids at 0, 48, and 96 hours after treatment with PPM as indicated. Monomeric Azami-Green1 (mAG1) is representative for S/G2/M-phase, mKusabira-Orange2 (mKO2) is representative for G1 and early S-phase.

In order to investigate the cell cycle changes induced by the PPM in more detail and with improved timely and spatial resolution, we applied the FUCCI (Fluorescence Ubiquitin Cell Cycle Indicator) reporter concept [23]. Briefly, in this reporter system the accumulation and proteolytic degradation of the two fluorescence-labelled cell cycle proteins Cdt1 and Geminin (involved in control of licensing of replication origins) is monitored. Consequently, cells with enrichment for Cdt1 – which is specific for G1-Phase and origin firing in early S-phase onset, show red fluorescence, while cells in S/G2/M-phase of the cell cycle express Geminin and are thus labelled by green fluorescence as schematically illustrated in figure S2B. We integrated an autonomously functional FUCCI reporter construct into the *Rosa26* locus of MC38 cells and tracked the complete cell cycle of individual cells with and without the addition of different PPM doses in real time by time-lapse fluorescence microscopy. We subsequently analyzed the individual cell cycle and morphology of all cells with appropriate software tools as described in the methods section. Overall, using this approach we detected a significant effect of PPM treatment on cell number, nucleus size, and distribution of cells in G1/S versus S/G2/M phase (Fig. S2C-F). These effects appeared moderate with PPM 1:200 dilution, but substantial with PPM 1:100 dilution treatment. In particular, at PPM 1:100 conditions, we observed complete stagnation of cell number over a period of 72 hours (Fig. S2C), which was associated with an excessive increase in cell nucleus size (Fig. S2D) and a strong accumulation of Cdt1-expressing (*i.e.* red fluorescent) cells at an early stage (approx. 6-24 hours post PPM treatment, Fig. S2E). Furthermore, real-time microscopy showed that the relative number of cells in G2/M decreased over time (Fig. S2F). Altogether, the live imaging analysis of MC38-FUCCI cells suggest that the PPM triggers strong accumulation of Cdt1 in late G1 presumably resulting in inappropriate origin firing, irregular DNA re-replication and eventually cell death, which best explains our findings obtained with conventional FACS analysis (compare to figure 1D).

In a next step, we aimed to confirm our findings in a 3D cell culture model better reflecting the situation of colon cancer growth *in vivo*. To this end, we generated 3D spheroids from MC38-FUCCI cells and determined their growth kinetics and FUCCI-Reporter fluorescence in presence or absence of the PPM for 96 h. Importantly, we detected a complete and statistically significant stagnation of spheroid growth 24 h after initiation of PPM treatment until termination of the experiment (Fig. 1E and Fig. S2G). This was associated with PPM-dependent spheroid deformation and strong Cdt1 accumulation (*i.e.* red fluorescence) over time, while control spheroids were rather characterized by an increase in green fluorescence indicating mitotic activity (Fig. 1F and Movie S1). In summary, our data suggests that PPM also mediates strong inhibitory effects on three-dimensional tumor-like structures by inappropriate replication initiation and subsequent cell death.

### Liver metastasis growth of MC38 cells is greatly enhanced by the PPM

To characterize the effect of the PPM *in vivo*, we decided to use the well-established MC38 liver metastasis model in syngeneic and immunocompetent C57BL/6J mice [19]. 14 days after injection of MC38 cells into the spleen, mice were gavaged once daily with the PPM for 7 consecutive days (Fig. S1A). Unexpectedly, this resulted in a massive and highly significant aggravation of liver metastasis burden in the PPM-treated animals (PPM group) compared to control mice (Ctrl) (Fig. 2A-C). While *in vivo* growth of MC38 cells after intrasplenic injection was restricted to the liver under control conditions, PPM treatment resulted in peritoneal dissemination of MC38 cells (Fig. S3A), further underscoring the enhanced malignancy after PPM gavage. Microscopically, well-demarcated nodular foci of atypical cells exhibiting mixed spindle and epithelioid morphology were identified in the livers. High mitotic activity with numerous mitotic and apoptotic figures was present. Multinucleated tumor giant cells were frequently observed. Heterogeneously distributed areas of necrosis comprised less than 5% of the total tumor volume. Focal lymphatic vascular invasion was noted. Focal areas of univacuolar adipose tissue were identified within the tumor lesion. The surrounding hepatic parenchyma demonstrated no significant histopathological alterations (Fig. S3B). Careful analysis of body weight over time and laboratory values suggested a good tolerability of the PPM in mice (Fig. S4). The significantly elevated transaminase AST in the PPM group (Fig. S4B) is most likely a consequence of peri-metastatic hepatocyte disintegration.

**Figure 2.**
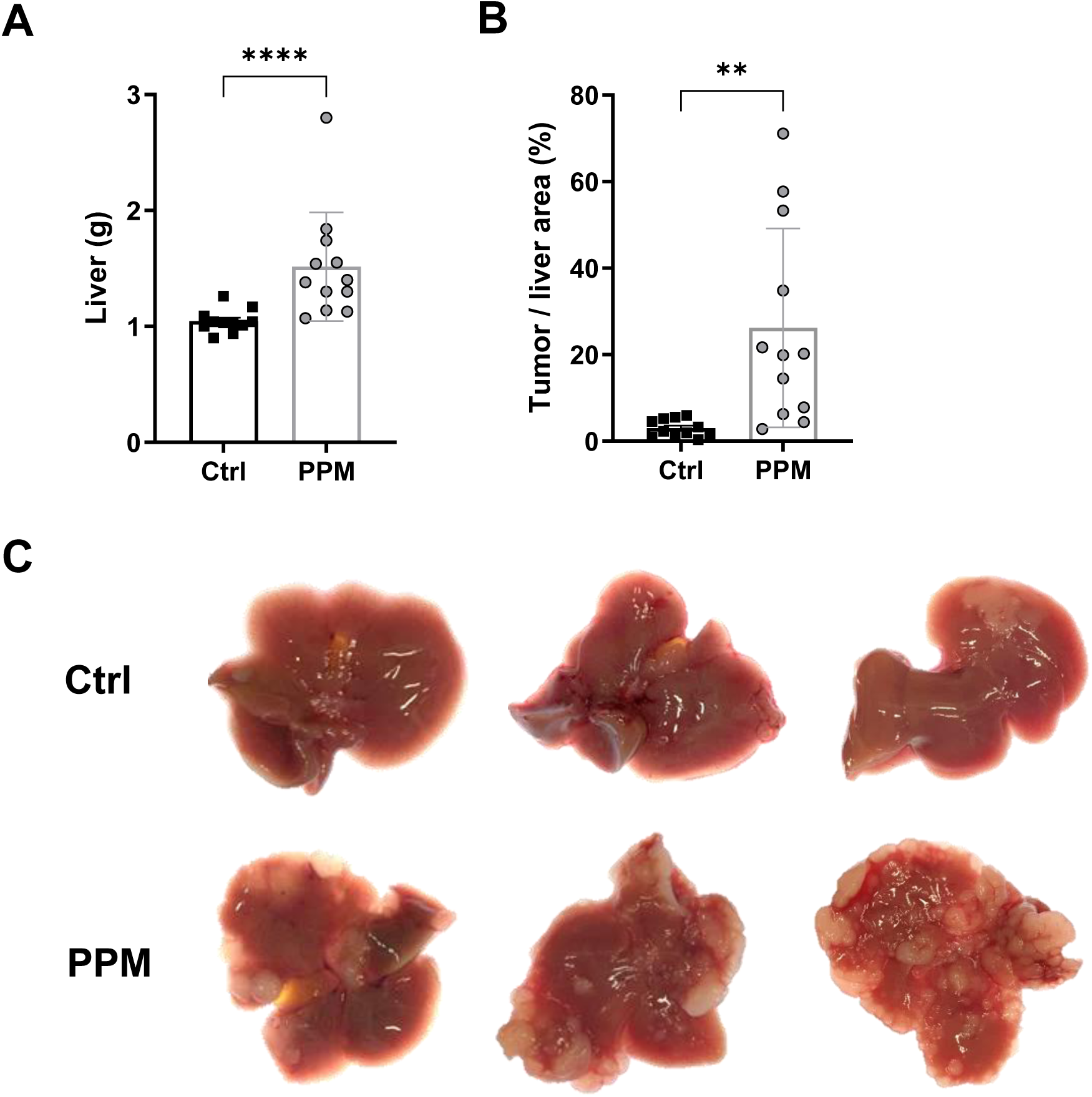
Effect of the PPM on MC38 liver metastases *in vivo*. **A.** Liver weight of Ctrl (*n* = 11) and PPM-treated (*n* = 12) groups at the end of the experiment. **B**. Tumor area was normalized to liver area and presented as percentage. Data represent means with SD and ** *p* ≤ 0.01, **** *p* ≤ 0.0001 according to Mann Whitney U test and Student’s unpaired t-test. **C.** Representative liver images from Ctrl and PPM-treated mice.

### PPM treatment results in significantly greater peri-metastatic immune cell abundance

Next, we sought to identify the mechanisms that underlie the metastasis-aggravating effect of the PPM *in vivo*. As inflammation is a well-established driver of liver metastasis [38], we decided to characterize the abundance of immune cells in our model. Total leukocytes (identified by immunohistology against CD45 (Fig. 3A)) and F4/80-positive macrophages (Fig. 3B) were evenly distributed in and around the MC38 liver metastases under control conditions. Oral application of the PPM led to significantly greater abundance of CD45-positive leukocytes (Fig. 3A) and, especially, F4/80-positive macrophages (Fig. 3B) in the peri-metastatic regions. These findings strongly support a situation where the PPM aggravates CRC liver metastasis through attraction of tumor-associated macrophages (TAMs) [39]. Next, we sought to better understand the molecular underpinnings of the observed TAM attraction. Gene expression analysis of several pro-inflammatory cyto- and chemokines via quantitative PCR revealed a significant upregulation of *Il-6* and *Ccl-2* in the liver (Fig. 4A). To further interrogate the cellular source for CCL-2, MC38 cells were stimulated *in vitro* with the PPM. As can be seen in figure 4B, the PPM significantly increased *Ccl-2* gene expression in MC38 cells, suggesting that the metastatic cells are one source for the chemokine CCL-2 in response to the PPM *in vivo*. A central importance of the CCL-2/TAM-axis in our project is supported by published data demonstrating an essential function of CCL-2 for the successful establishment of MC38 liver metastases [19]. Together, these data show PPM-induced inflammation around the metastatic nodules and argue for a causal role of the chemokine CCL-2 for the attraction of TAMs to metastatic nodules.

**Figure 3.**
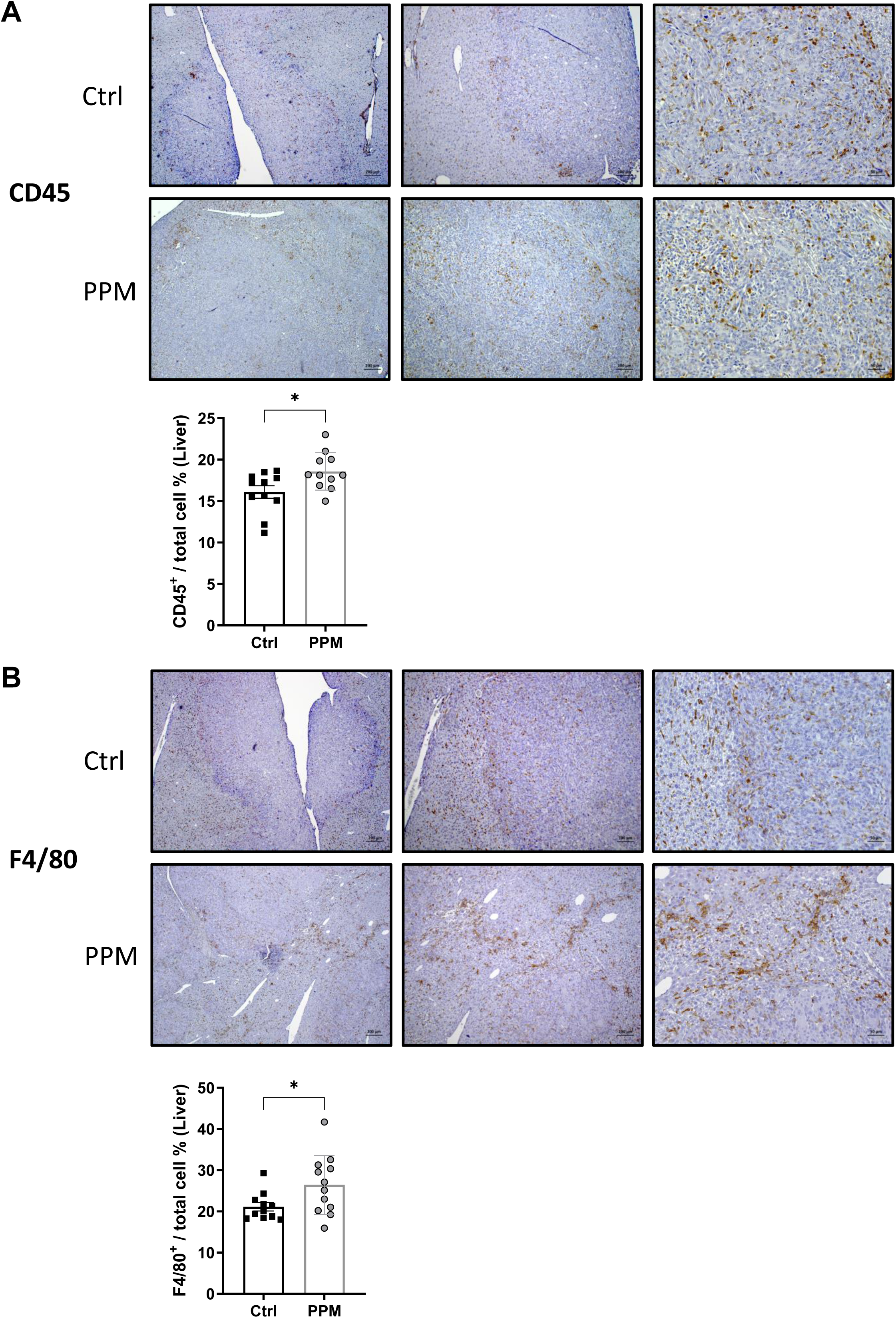
Representative images of CD45 (**A**), and F4/80 (**B**) IHC staining in liver tissues from Ctrl and PPM-treated groups. The images were taken at magnifications of 50x, 100x, and 200x from left to right (scale bars are 200, 100, and 50 µm, respectively). Graphs show the percentage of positive cells in the liver areas, detected on IHC staining images using QuPath software. Data represent means with SD, * *p* < 0.05 according to Student’s unpaired t-test (A) and Mann Whitney U test (B).

**Figure 4.**
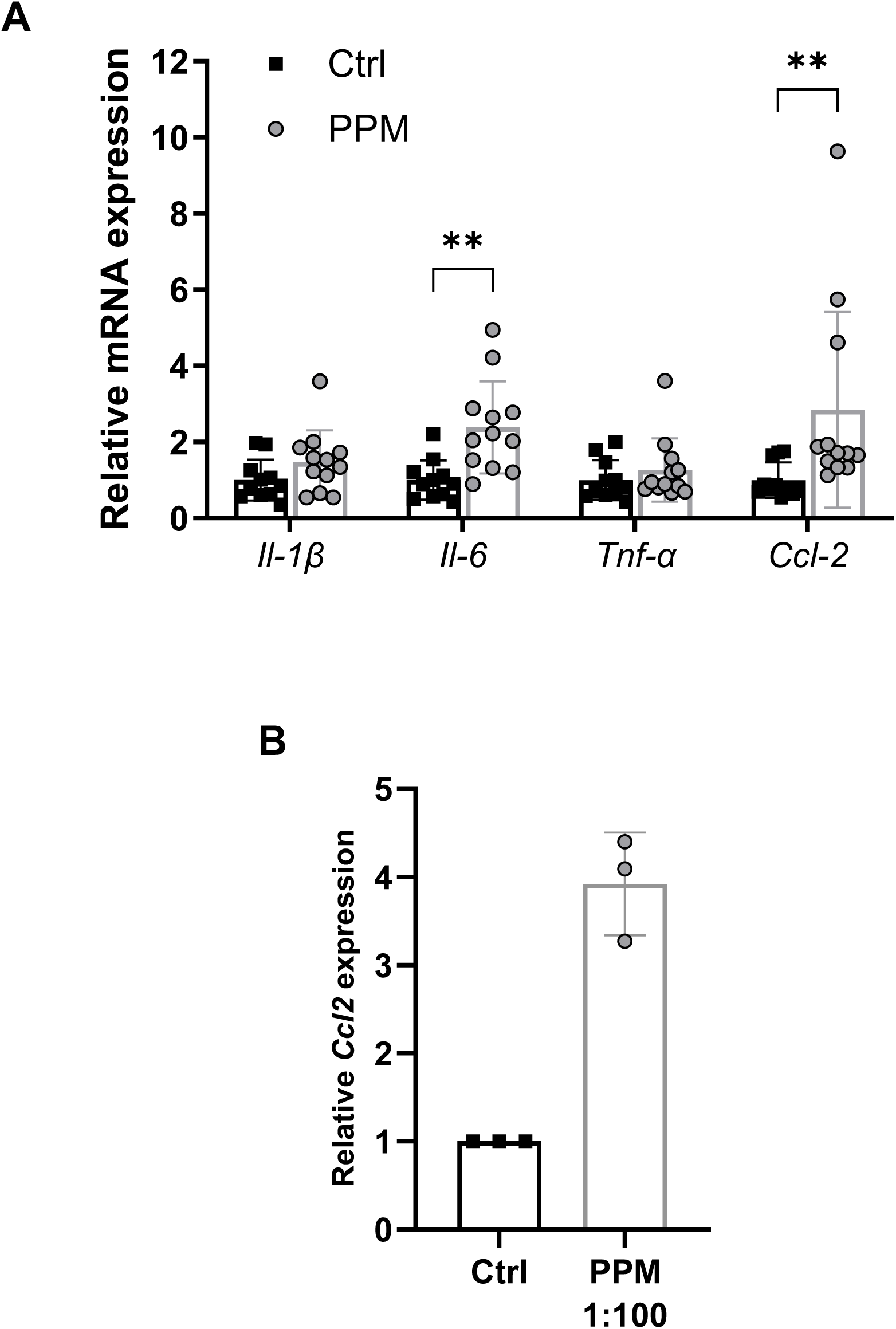
A. Effect of the PPM on gene expression of pro-inflammatory cyto- and chemokines in murine liver harboring MC38 metastases. **B.** Gene expression of *Ccl2* in Ctrl and PPM (1:100 dilution)-treated MC38 cells at 48h in cell culture. Data represent the means with SD and ** *p* ≤ 0.01 according to the Student’s unpaired t-test or Mann Whitney U test analysis (A).

### Functional importance of inflammation for PPM-induced growth of liver metastases

To test the hypothesis that peri-tumoral inflammation underlies the metastasis-promoting effect of the PPM, experiments with the broad-spectrum anti-inflammatory agent prednisolone [26] were performed (Fig. S1B). Strikingly, the enhancing effect of the PPM on liver weight and relative number of metastatic nodules observed before (see Fig. 2) was completely blunted by co-administration of prednisolone (Fig. 5). Taken together, these results strongly support the hypothesis that enhanced inflammatory activity underlies the metastasis-promoting effect of the PPM in our model. Tolerability analyses did not result in safety concerns of the combined application of the PPM and prednisolone (Fig. S5). As glucose is an important fuel for cancer cells and elevated blood glucose is a well-established side effect of prednisolone, we quantified blood glucose values in this setting (Fig. S5C). No differences between the experimental groups were noted, strongly arguing against a potential confounding effect of hyperglycemia after prednisolone.

**Figure 5.**
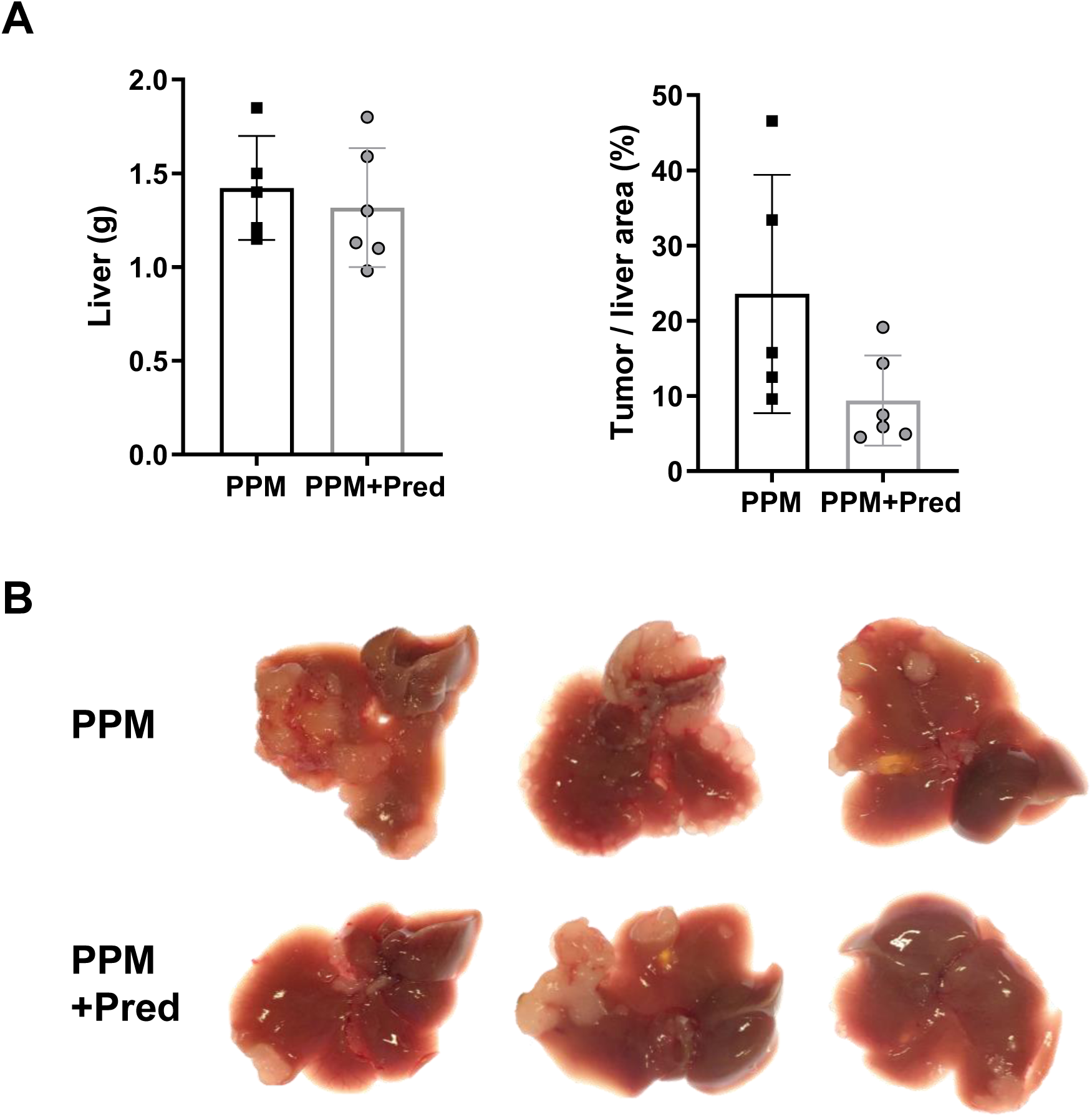
Importance of inflammation for PPM-induced growth of liver metastasis. **A.** Liver weight and tumor area calculation of PPM (*n* = 5) and PPM-treated (*n* = 6) groups. Tumor area was normalized to liver area and presented as percentage. The images were captured at magnifications of 50x, 100x, and 200x from left to right (scale bars are 200, 100, and 50 µm, respectively). **B.** Representative liver images from mice treated with PPM and those additionally injected with prednisolone (PPM+Pred).

### The pro-inflammatory effect of the PPM depends on the existence of colon cancer cells in the liver

To further investigate the nature of the inflammation-inducing signal, metastasis-free mice were gavaged with the PPM (Fig. S1C). This approach did not result in detectable changes, neither in the blood nor in the liver of the mice (Fig. S6). These results strongly argued against a functional significance of liver cells (e.g. hepatocytes, cholangiocytes or hepatic stromal cells) for the inflammatory reaction caused by the PPM when the liver is devoid of tumor cells. Furthermore, these results clearly demonstrated that the applied dose of the PPM was non-toxic to the mice.

### Effect of the PPM on the intestinal microbiota of mice with MC38 liver metastases

The intestinal microbiota is causally linked to local growth and metastasis of colon cancer [40–43]. In addition, consumption of various plant polyphenols has been shown to influence gut microbial communities and function [44, 45]. Based on this, we analyzed the changes in the cecal microbiota of mice for the metastasis-aggravating effect of the PPM, using 16S rRNA gene amplicon sequencing. To assess the effects of PPM itself on the microbiota, we included also PPM-treated metastasis-free (healthy) mice. PPM treatment did not result in a significant change in α-diversity based on both Shannon diversity index (Fig. 6A) and richness (Fig. S7A) for mice with CRC liver metastasis (Ctrl and PPM) and metastasis-free mice (depicted as Ctrl (-LM) and PPM (-LM)). For CRC liver metastasis mice, microbiota profiles between individuals (β-diversity) were also not influenced by PPM treatment (Fig. 6B, Fig. S7B, C). However, the only effect of PPM was observed in metastasis-free mice showing increased *ASF356* genus in PPM (-LM) compared to Ctrl (-LM) in all the tested taxa (Fig. 6C). Interestingly, for Ctrl vs Ctrl (-LM) and PPM vs PPM (-LM) comparisons, a significant shift in the gut microbiota was observed in response to the surgical procedure for tumor cell injection, based on the two β-diversity indexes tested (Bray-Curtis and Jaccard dissimilarity indices, Fig. 6B, Fig. S7B and C). The mice that underwent surgery (Ctrl group) had less *Lactobacillaceae* based on relative abundances, with underlying changes at the genus level: decrease of *HT002*, *Dubosiella* and increase of *Faecalibaculum*, *ASF356*, [*Eubacterium]_xylanophilum_group* compared to Ctrl (-LM) group (Fig. 6C).

**Figure 6.**
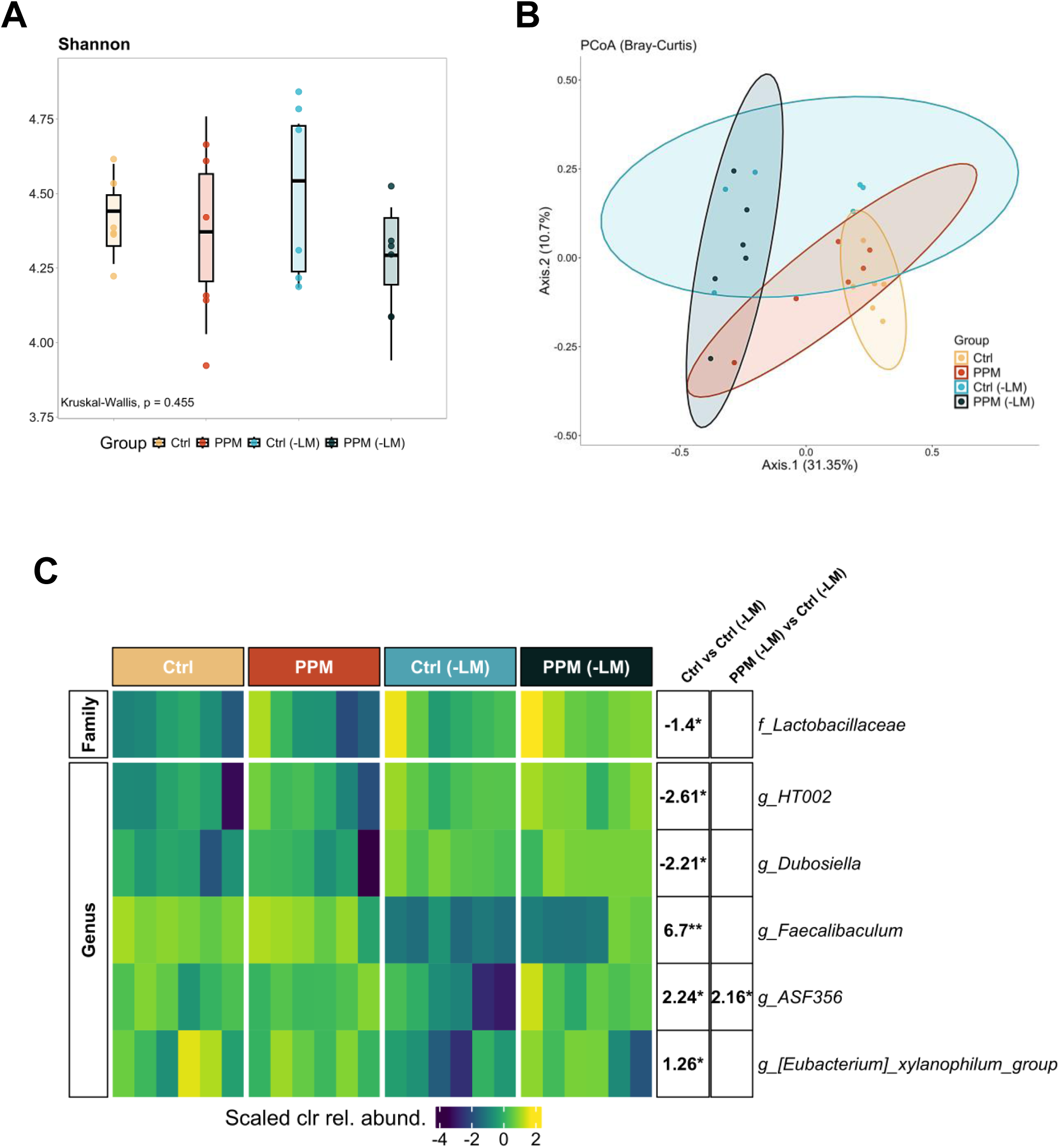
A. Alpha diversity analysis of the four groups using Shannon diversity index. Each sample was represented by points, and the boxplots show the interquartile ranges (IQRs) of the data covered by boxes, median values shown by the horizontal black line, and ± 1.5-fold the IQR represented by whiskers. Statistical comparison of the groups was conducted using Kruskal-Wallis test and the obtained *p* value was shown on the figure. **B.** Beta diversity analysis based on Bray Curtis dissimilarities. Principal coordinates analysis (PCoA) plot was produced using the first two axes which represent 31.35% and 10.7% of the variation of the data, respectively. Points represent sample scores, and the ellipses correspond to 95% confidence regions for mouse groups. **C.** Taxa at the family and genus levels were detected to be significantly different from Ctrl vs Ctrl (-LM) and PPM (-LM) vs Ctrl (-LM) comparisons. Scaled centered-log-ratio (clr) transformed relative abundance values of each taxon are shown in the heatmap for each group. Log2 fold change values with significance stars representing adjusted *p* values from Kruskal Wallis followed by Dunn’s test with Benjamini-Hochberg correction are given in the right annotation columns for respective pairwise comparisons of each taxon (* 0.01 < *p*.adj < 0.05; ** 0.001 < *p*.adj ≤ 0.01; *** *p*.adj ≤ 0.001) (Ctrl, *n* = 6; PPM, *n* = 6; Ctrl (-LM), *n* = 6; PPM (-LM), *n* = 6).

**Figure 7.**
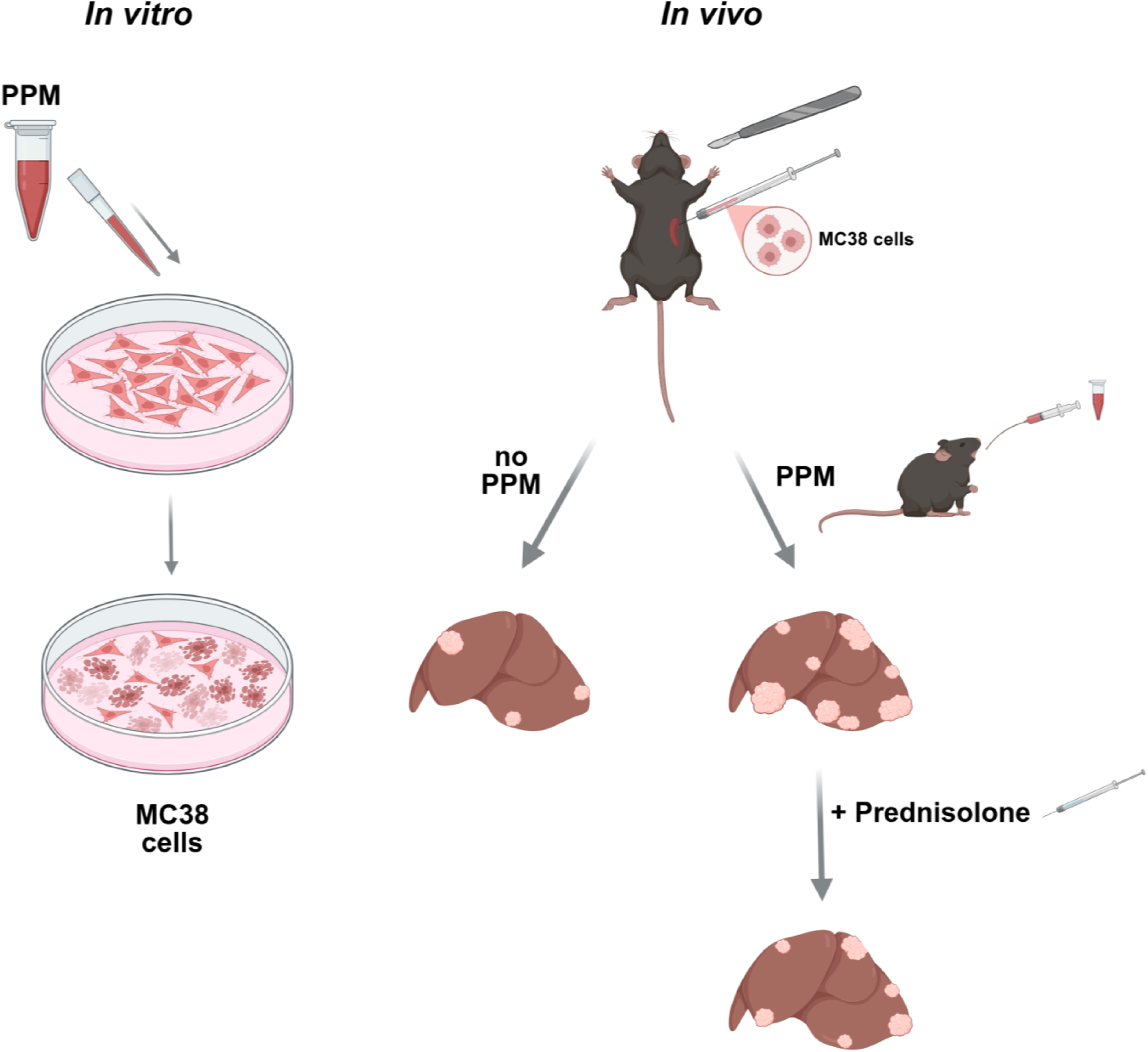
Graphical summary of the results.

## Discussion

Our pre-clinical study demonstrated substantial differences regarding the effects of a mixture of different plant polyphenols on the growth and survival of the established murine colon cancer cell line MC38 *in vitro* versus *in vivo*. While the plant polyphenol mix (PPM) used by us strongly inhibited survival, proliferation and cell cycle progression of the cells in the culture dish, it’s application via oral gavage to mice harboring MC38 liver metastases significantly aggravated malignant progression. We present experimental evidence to support a causal role of enhanced inflammatory activity for the metastasis-boosting effect of the PPM. While our *in vitro* results are well in line with the published literature, the observation that plant polyphenols can accelerate metastasis formation *in vivo* is new and scrutinizes the notion that plant polyphenols should be investigated as substances for cancer therapy.

In the present study we provided evidence that PPM treatment of MC38 colon carcinoma cells results in aberrant cell cycle progression, strongly increased DNA synthesis and cell death. In conventional FACS analyses this phenomenon was associated with cell populations showing DNA contents <2n (“sub-G1 peak”), which is typical for apoptotic cells, as well as a large population of cells with a DNA content >4n referred to as “supra-G2” cells. More detailed time lapse analysis using a FUCCI cell cycle Reporter integrated into MC38 cells revealed that PPM triggers stabilization of the Cdt1 protein and enhanced degradation of its inhibitor Geminin. Cdt1 is an important component of the pre-replication complex and thus relevant for precise single origin firing thereby preventing DNA re-replication [46]. However, inappropriate increase of Cdt1 in late S and early G2 poses a potential risk of replication origin over-firing and re-replication, which could occur if there were residual activity of the DNA-replicating enzymes in G2. The activity of Cdt1 is tightly controlled by Geminin, a re-replication inhibitory factor, which directly binds to and suppresses the replication-stimulating function of Cdt1 [46]. We conclude that PPM-driven Cdt1 accumulation may also result on over-firing at replication origins eventually resulting in a phenomenon referred to as replication catastrophe as reviewed by Toledo *et al.* [47], which best explains the phenotype, Cdt1 expression pattern and morphology of PPM-dependent MC38-FUCCI cells. However, the signaling mechanism linking PPM to Cdt1 stabilization needs to be clarified in future studies.

The scientific literature is replete with studies that report “anti-tumor” activity of plant-derived molecules against established cancer cell lines [9, 48]. Various pro-tumorigenic pathways can be affected by plant polyphenols, e.g. proliferation, growth-promoting signal transduction pathways, resistance to apoptosis, induction of angiogenesis, migration and invasion, epithelial-to-mesenchymal transition, cellular energetics and resistance to therapy, resulting in the notion that phytochemicals exert “pleiotropic anti-cancer effects” [49]. Our *in vitro* data are well in line with this and show significant and dose-dependent inhibitory effects of the used PPM on survival, proliferation and cell cycle progression of MC38 cells. The most striking observation was the appearance of large cells with greatly increased DNA content in the supra-G2 fraction of the cell cycle after treatment with the PPM. These cells resemble “polyploid giant cancer cells” (PGCC), which can be found in both rodent tumor models as well as human cancers under conditions of cellular stress (e.g. nutrient depletion, hypoxia, radiation, chemotherapy) [37]. Our finding of large polyploid cells is in line with published literature describing inhibitory effects of plant polyphenols on cell cycle progression and cytokinesis [50–52]. We were intrigued to observe the formation of giant cancer cells not only *in vitro*, but also in the liver metastases in response to oral application of the PPM. This underscores the notion that the PPM has significant inhibitory effects on cell cycle progression and strongly argues for sufficient bioavailability and *in vivo* functionality of our polyphenol mixture. Whether the giant cells are responsible for the aggravated inflammatory response seen *in vivo* via enhanced expression and secretion of *Ccl-2* is an intriguing and important question that lies out of the scope of the current project but should be functionally interrogated in future studies.

While a vast amount of original studies and reviews on the role of phytochemicals for colon cancer have been published [53, 54], the effects of these substances for liver metastasis of colon cancer are less well understood. Narayanan *et al.* investigated an extract of seeds from the melinjo tree (the active polyphenol is called gnetin C) and observed a dose-dependent inhibition of liver metastases of the established murine CRC cell line colon-26 in syngeneic BALB/c mice [55]. Yuan *et al.* analyzed the effect of epigallocatechin gallate (EGCG, one of the many different polyphenols found in green tea) on CRC liver metastasis with xenotransplants of human HT-29 cells in BALB/c nude mice. Again, statistically significant and dose-dependent inhibitory effects were observed and the highest dose even completely abrogated liver metastasis formation [56]. On the contrary, a mixture of bioactive compounds from the *Ginkgo biloba* tree strongly aggravated CRC liver metastasis formation of xenotransplanted human SW-620 cells in athymic nude mice [57]. Our observation of significantly aggravated hepatic metastasis by plant polyphenols is in line with the latter study. However, the mechanistic underpinnings of the metastasis-inducing effect reported by Wang et al. [57] remain elusive as the authors did not test the functional relevance of the observed changes in gene expression and MAP kinase activity. Taken together, both inhibitory and exacerbating effects of plant polyphenols on rodent models of CRC liver metastases have been published, arguing for additional studies into this topic.

Most studies regarding the effects of phytochemicals on tumor progression have used immunodeficient mice, most prominently xenografts of established human cancer cell lines. This experimental approach completely negates the central importance of the immune system for cancer progression [58] and does not allow to characterize how phytochemicals impact on the activity of immune cells in the context of cancer. This is especially relevant as plant polyphenols are typically regarded as having robust anti-inflammatory effects [59]. Our results stand in contrast with this notion and demonstrate clear pro-inflammatory effects of plant polyphenols both *in vitro* and *in vivo*. We provided experimental evidence for a causal role of this pro-inflammatory action for the observed metastasis-aggravating effect of the plant polyphenol mixture used by us. It has been known for a long time that cancer cells are able to release chemotactic factors (e.g. CCL2, SDF-1α and mCSF1) to attract pro-tumorigenic immune cells, e.g. monocytes and neutrophils, to tumors [60–62]. Our results are well in line with these data and add phytochemicals to the list of inducers of inflammation such as immunogenic cell death, hypoxia, nutrient depletion and chemo/radiotherapy. Taken together, our data illustrate that the ultimate effect of a given intervention on cancer depends on a complex interaction of the cancer cells with multiple components of the host (e.g. the immune system) and cannot be determined with sufficient accuracy by in vitro experiments. The search term “phytochemicals and cancer” results in roughly 11.000 hits in PubMed and since the year 2021, over 1.000 studies fitting to these keywords have been published annually. These studies almost exclusively analyze the effects of plant polyphenols and other phytochemicals on established cancer cell lines *in vitro*. Our results underscore the notion that results from *in vitro* studies and xenograft models should be thoroughly scrutinized with model systems that more closely resemble human cancer before a true tumor-inhibiting effect can be assumed [63]. This argues for critical appraisal of experiments that report only cell culture data and once again underscores the central importance of the used rodent model (immunodeficient vs. immunocompetent).

## Author Contributions

Conceptualization M.E. and T.Cr.; Experimentation J.R., J.K., J.N., H.H. and M.E.; Data Analysis and Visualization J.R., J.K., J.N., E.S., F.S., N.T., T.Cl., H.H., C.L., M.E. and T.Cr.; Writing M.E., H.H., C.L. and T.Cr.; Supervision, Resources, and Funding Acquisition T.Cr.; Project Administration M.E. and T.Cr.

## Acknowledgements

We are grateful to Ntana Kousetzi for her technical assistance with sample processing for 16S rRNA gene amplicon analysis. Research in the Cramer lab was supported by grants from the Deutsche Forschungsgemeinschaft (CR 133/2-1 until 2-4 and CR 133/3-1). Christian Liedtke is funded by the German Research Foundation (DFG), CRC1382-403224013, project A02. Thomas Clavel received funding from the DFG, project ID 403224013, CRC1382 project Q02. This work was supported by the Flow Cytometry Facility of the Interdisciplinary Center for Clinical Research (IZKF) within the Faculty of Medicine at RWTH Aachen University.

## Supplementary Online Materials

**Table S1.** Composition of the plant polyphenol mixture:

| **Ingredient** | **Amount** |
| --- | --- |
| Beetroot | 100 g |
| Broccoli | 100 g |
| Brussels sprouts | 100 g |
| Cauliflower | 100 g |
| Spinach | 100 g |
| Lemon zest | 12.5 g |
| Long pepper (Piper longum) | 6 g |
| Garlic | 100 g |
| Turmeric powder | 8 g |
| Linseed oil | 10 ml |
| Green tea polyphenols (LKT Labs G6817) | 2.4 g |

## Supplementary Figure Legends

**Figure S1** Schematic representation of the experimental setups used in this study. **A.** 14 days after MC38 cell injection and splenectomy, mice were treated with either PPM for seven days (showed as PPM group in the figures) or left untreated (Ctrl group). **B.** 14 days after MC38 cell injection and splenectomy, mice were treated with PPM (PPM group) and injected with prednisolone (PPM+Pred group) for seven days. **C.** Metastasis-free (healthy) mice were treated with either PPM for seven days (PPM (-LM) group) or left untreated (Ctrl (-LM) group).

**Figure S2 A.** Cell cycle analysis of Ctrl and PPM (at 1:100 dilution)-treated MC38 cells using flow cytometry and propidium iodide staining. The relative cell number percentages of different cell cycle phases were calculated from the plots in Figure 1D and presented in the graph. Relative cell number distribution of different cell cycle phases after flow cytometry analysis of propidium iodide-stained MC38 cells after 48 hours under control conditions (Ctrl) or PPM treatment in 1:100 dilution. **B.** Basic principle of Fluorescence Ubiquitin Cell Cycle Indicator (FUCCI) technology. Cells expressing the FUCCI reporter system will either fluoresce in green when in late S, G2 or M-phase and red when in G1 until early S-phase. **C-F.** MC38-FUCCI cells were cultivated either in absence or presence of PPM at concentrations indicated over the course of 72 hours in the incubation chamber of an Axio Observer.Z1/7 microscope und subjected to time-lapse imaging for brightfield, green (Alexa 488) and red (Alexa XYZ) fluorescence3, respectively. Live cell image analysis data are displayed as mean of *n* = 16 measurements per condition every 15 minutes over 72 hours; bars indicating SD were omitted for visual clarity. Relative cell number (**C**), relative nucleus size (**D**), relative cell number of cells in G1/S-phase (**E**), and relative cell number of cells in S/G2/M-phase (**F**). Relative numbers were normalized to respective baseline measurements of each treatment condition at 0 hours. **G.** Complete group comparisons of Figure 2E were presented on the table. Statistical significances in C-G were determined using Dunnett’s multiple comparisons test, comparing PPM treatments to control (\*\**p* < 0.01, \*\*\**p* < 0.001, \*\*\*\**p* < 0.0001).

**Figure S3 A.** Pictures of PPM-treated mice show the dissemination of MC38-induced tumors on the peritoneal wall. **B.** Representative images of hematoxylin and eosin (H&E) staining in liver tissues from Ctrl and PPM-treated groups. The images were taken at magnifications of 50x, 100x, and 200x from left to right (scale bars are 200, 100, and 50 µm, respectively).

**Figure S4: A.** Body weight follow-up and body weight change of Ctrl (*n* = 5) and PPM (*n* = 6) mice during the experiment period. **B.** Serum markers of liver toxicity, **C.** Measurement of complete blood count after the sacrifice of Ctrl (*n* = 11) and PPM (*n* = 12) mice groups. Data represent means with SEM (**A**) or SD (**B**, **C**) and ** *p* ≤ 0.01 according to two-way analysis of variance (ANOVA) with Šídák’s multiple comparisons test (**A**), Student’s unpaired t-test (**B**) and Mann Whitney U test (**C**).

**Figure S5: A.** Body weight follow-up and body weight change of mice treated with PPM (*n* = 6) and those additionally injected with prednisolone (PPM+Pred) (*n* = 6). **B.** Serum markers of liver toxicity, **C.** Blood glucose, **D.** Measurement of complete blood count after the sacrifice of PPM (*n* = 5) and PPM+Pred (*n* = 5-6) mice groups. Data represent means with SEM (**A**) or SD (**B**, **C**, **D**) and were analyzed by two-way analysis of variance (ANOVA) with Šídák’s multiple comparisons test (**A**), Student’s unpaired t-test (**B**) and Mann Whitney U test (**C, D**).

**Figure S6 A.** Body weight follow-up and body weight change of metastasis-free mice groups, either untreated, (Ctrl (-LM), *n* = 6) or PPM-treated (PPM (-LM), *n* = 6) groups. Liver weight (**B**), serum markers of liver toxicity (**C**) and measurement of complete blood count (**D**) after the sacrifice of Ctrl (-LM) (*n* = 6) and PPM (-LM) (*n* = 5-6) groups. Data represent means with SEM (**A**) or SD (**B**, **C**, **D**) and were analyzed by two-way analysis of variance (ANOVA) with Šídák’s multiple comparisons test (**A**), Student’s unpaired t-test (**B**, **C**) and Mann Whitney U test (**D**).

**Figure S7 A.** Alpha diversity analysis based on observed ASV numbers. Each sample was represented by points, and the boxplots show the interquartile ranges (IQRs) of the data covered by boxes, median values shown by the horizontal black line, and ± 1.5-fold the IQR represented by whiskers. Statistical comparison of the groups was conducted using Kruskal-Wallis test and the obtained *p* value was shown on the figure. For pairwise comparisons of the groups, post-hoc Dunn’s Test with Benjamini-Hochberg correction was performed. NS represents no significance (*p*.adj > 0.05). **B.** Pairwise Permutational Multivariate Analysis of Variance (PERMANOVA) test results for the beta diversity analysis based on Bray-Curtis dissimilarities. **C.** Beta diversity analysis based on Jaccard index. Principal coordinates analysis (PCoA) plot was produced using the first two axes which represent 21.49% and 8.7% of the variation of the data, respectively. Points represent sample scores and the ellipses correspond to 95% confidence regions for mouse groups. Pairwise PERMANOVA test results were submitted in the table on the right side (* 0.01 < *p*.adj < 0.05; ** 0.001 < *p*.adj ≤ 0.01; *** *p*.adj ≤ 0.001) (Ctrl, *n* = 6; PPM, *n* = 6; Ctrl (-LM), *n* = 6; PPM (-LM), *n* = 6).

## Supplementary Methods

### Primer sequences used in the gene expression analyses

The primer sequences of each gene used in quantitative real-time PCR experiments were as follows: *B2m* (F: TTCTGGTGCTTGTCTCACTGA, R: CAGTATGTTCGGCTTCCCATTC), *Il-1β* (F: CAACCAACAAGTGATATTCTCCATG, R: GATCCACACTCTCCAGCTGCA), *Il-6* (F: TGAGAAAAGAGTTGTGCAATGGC, R: GCATCCATCATTTCTTTGTATCTCTGG), *Tnf-α* (F: CCATTCCTGAGTTCTGCAAAGG, R: AGGTAGGAAGGCCTGAGATCTTATC), *Ccl-2* (F: GCTGTAGTTTTTGTCACCAAGC, R: GACCTTAGGGCAGATGCAGT), *Ho-1* (F: AAGCCGAGAATGCTGAGTTCA, R: GCCGTGTAGATATGGTACAAGGA), *Nqo-1* (F: AGAGAGTGCTCGTAGCAGGAT, R: CTACCCCCAGTGGTGATAGAAA).

### Bioinformatic Analysis of 16S rRNA Gene Amplicon Sequencing Data

Quality controls of the reads were carried out using FastQC (ver. 0.12.1), and trimming was performed using fastp (ver. 0.23.4) [1] with parameter settings of “qualified_quality_phred” = 20 and “length_required” = 200. Then, quality trimmed reads were imported to Qiime2 (ver. 2023.09) [2]. Primer sequences (Forward: 5′-CCTACGGGNGGCWGCAG-3′, Reverse: 5′-GACTACHVGGGTATCTAATCC-3′) were removed using Cutadapt plugin [3]. DADA2 pipeline was employed for denoising and construction of ASVs (i.e. amplicon sequence variants) [4]. Since DADA2 produces an error model for each sequencing run, the steps above were separately followed for two different run batches, and denoised data were then merged before downstream procedure. For production of phylogenetic tree, ‘q2-phylogeny’ plugin was used. For the taxonomic classification of the ASVs, a mouse stool weighted taxonomic classifier was created by processing SILVA 138.1 SSURef NR99 full-length reads [5] to extract amplicon region-specific reads matching with the primer sequences above, dereplicating using ‘RESCRIPt’ plugin [6], and finally using a Scikit-learn naïve Bayes machine-learning taxonomy classifier with mouse stool weights which were assembled using ‘q2-clawback’ plugin. Rotted phylogenetic tree, ASV feature abundance matrix, and taxonomic information of the ASV features were exported. All downstream data analyses were carried out in R (ver. 4.2.2).

### Diversity Analysis

Using ASV count and taxonomy tables along with metadata, a phyloseq object was created using phyloseq package (ver. 1.52.0) [7]. For alpha diversity analysis, rarefication was performed using the minimum sample size of 4249. Three indices; observed, Shannon and Simpson were calculated by “estimate_richness” function of phyloseq package, and Kruskal-Wallis (KW) and post-hoc Dunn’s test with Benjamini-Hochberg (BH) correction were performed to test significance of the changes between mice groups, and adjusted *p* value (*p*.adj) of 0.05 was used as a significance threshold. For beta diversity analysis, non-rarefied count matrices were used, and Bray-Curtis and Jaccard indexes were calculated. PCoA plots were prepared using first two axes. For significant testing, Permutational multivariate analysis of variance (PERMANOVA) was used via “adonis2” function in vegan package (ver. 2.7.1) [8]. For pairwise comparisons, “pairwise.adonis2” function in pairwiseAdonis package (ver. 0.4.1) [9] was used.

### Differential Relative Abundance Analysis

For taxonomic differential abundance (DA) analysis, ASV counts were aggregated into phylum, family and genus levels. Unclassified features at each level were removed. For each level, rare taxa were filtered out before DA if any taxa not detected in at least 90% of the samples in each group with at least 0.1% relative abundance for phylum or 0.25% for family and genus levels [10]. KW and post-hoc Dunn’s test with BH correction were carried out, and p.adj threshold was set to 0.05. For heatmaps, centered-log-ratio values were calculated, and scaling was performed. Heatmaps were obtained using ComplexHeatmap (ver. 2.24.1) package [11].

## References

1. Mattiuzzi C, Sanchis-Gomar F, Lippi G. Concise update on colorectal cancer epidemiology. Ann Transl Med. 2019;7(21):609. Epub 2020/02/13. doi: 10.21037/atm.2019.07.91. PubMed PMID: 32047770; PubMed Central PMCID: PMCPMC7011596.

2. Siegel RL, Miller KD, Fedewa SA, Ahnen DJ, Meester RGS, Barzi A, et al. Colorectal cancer statistics, 2017. CA: A Cancer Journal for Clinicians. 2017;67(3):177–93. doi: 10.3322/caac.21395.

3. Zarour LR, Anand S, Billingsley KG, Bisson WH, Cercek A, Clarke MF, et al. Colorectal Cancer Liver Metastasis: Evolving Paradigms and Future Directions. Cellular and Molecular Gastroenterology and Hepatology. 2017;3(2):163–73. doi: 10.1016/j.jcmgh.2017.01.006.

4. Dhir M, Sasson AR. Surgical Management of Liver Metastases From Colorectal Cancer. Journal of Oncology Practice. 2016;12(1):33–9. doi: 10.1200/JOP.2015.009407.

5. Fraser GE, Butler FM, Shavlik DJ, Mathew RO, Oh J, Sirirat R, et al. Longitudinal associations between vegetarian dietary habits and site-specific cancers in the Adventist Health Study-2 North American cohort. The American Journal of Clinical Nutrition. 2025;122(2):535–43. doi: 10.1016/j.ajcnut.2025.06.006.

6. Watling CZ, Schmidt JA, Dunneram Y, Tong TYN, Kelly RK, Knuppel A, et al. Risk of cancer in regular and low meat-eaters, fish-eaters, and vegetarians: a prospective analysis of UK Biobank participants. BMC Med. 2022;20(1):73. Epub 20220224. doi: 10.1186/s12916-022-02256-w. PubMed PMID: 35197066; PubMed Central PMCID: PMCPMC8867885.

7. Sharma E, Attri DC, Sati P, Dhyani P, Szopa A, Sharifi-Rad J, et al. Recent updates on anticancer mechanisms of polyphenols. Front Cell Dev Biol. 2022;10:1005910. Epub 20220929. doi: 10.3389/fcell.2022.1005910. PubMed PMID: 36247004; PubMed Central PMCID: PMCPMC9557130.

8. Rudzińska A, Juchaniuk P, Oberda J, Wiśniewska J, Wojdan W, Szklener K, et al. Phytochemicals in Cancer Treatment and Cancer Prevention-Review on Epidemiological Data and Clinical Trials. Nutrients. 2023;15(8). Epub 20230414. doi: 10.3390/nu15081896. PubMed PMID: 37111115; PubMed Central PMCID: PMCPMC10144429.

9. Cháirez-Ramírez MH, de la Cruz-López KG, García-Carrancá A. Polyphenols as Antitumor Agents Targeting Key Players in Cancer-Driving Signaling Pathways. Front Pharmacol. 2021;12:710304. Epub 20211020. doi: 10.3389/fphar.2021.710304. PubMed PMID: 34744708; PubMed Central PMCID: PMCPMC8565650.

10. Droguett D, Castillo C, Leiva E, Theoduloz C, Schmeda-Hirschmann G, Kemmerling U. Efficacy of quercetin against chemically induced murine oral squamous cell carcinoma. Oncol Lett. 2015;10(4):2432–8. Epub 20150812. doi: 10.3892/ol.2015.3598. PubMed PMID: 26622865; PubMed Central PMCID: PMCPMC4580003.

11. Tsouh Fokou PV, Kamdem Pone B, Appiah-Oppong R, Ngouana V, Bakarnga-Via I, Ntieche Woutouoba D, et al. An Update on Antitumor Efficacy of Catechins: From Molecular Mechanisms to Clinical Applications. Food Science & Nutrition. 2025;13(4):e70169. doi: 10.1002/fsn3.70169.

12. Bimonte S, Barbieri A, Palma G, Luciano A, Rea D, Arra C. Curcumin inhibits tumor growth and angiogenesis in an orthotopic mouse model of human pancreatic cancer. Biomed Res Int. 2013;2013:810423. Epub 20131110. doi: 10.1155/2013/810423. PubMed PMID: 24324975; PubMed Central PMCID: PMCPMC3842048.

13. Chen L, Musa AE. Boosting immune system against cancer by resveratrol. Phytother Res. 2021;35(10):5514–26. Epub 20210608. doi: 10.1002/ptr.7189. PubMed PMID: 34101276.

14. Castillo-Pichardo L, Martínez-Montemayor MM, Martínez JE, Wall KM, Cubano LA, Dharmawardhane S. Inhibition of mammary tumor growth and metastases to bone and liver by dietary grape polyphenols. Clin Exp Metastasis. 2009;26(6):505–16. Epub 20090318. doi: 10.1007/s10585-009-9250-2. PubMed PMID: 19294520; PubMed Central PMCID: PMCPMC2898569.

15. Gingras D, Béliveau R. TOWARDS A GLOBAL ASSESSMENT OF THE ANTICANCER PROPERTIES OF FRUITS AND VEGETABLES: THE MONTREAL ANTICANCER NUTRINOME PROJECT. Acta Horticulturae. 2007:157–64. doi: 10.17660/ActaHortic.2007.744.15.

16. Sarkar FH, Li Y. Harnessing the fruits of nature for the development of multi-targeted cancer therapeutics. Cancer Treat Rev. 2009;35(7):597–607. Epub 20090805. doi: 10.1016/j.ctrv.2009.07.001. PubMed PMID: 19660870; PubMed Central PMCID: PMCPMC2784186.

17. Bock C, Lengauer T. Managing drug resistance in cancer: lessons from HIV therapy. Nature Reviews Cancer. 2012;12(7):494–501. doi: 10.1038/nrc3297.

18. Mai V, Colbert LH, Berrigan D, Perkins SN, Pfeiffer R, Lavigne JA, et al. Calorie restriction and diet composition modulate spontaneous intestinal tumorigenesis in Apc(Min) mice through different mechanisms. Cancer Res. 2003;63(8):1752–5. PubMed PMID: 12702556.

19. Zhao L, Lim SY, Gordon-Weeks AN, Tapmeier TT, Im JH, Cao Y, et al. Recruitment of a myeloid cell subset (CD11b/Gr1 mid) via CCL2/CCR2 promotes the development of colorectal cancer liver metastasis. Hepatology. 2013;57(2):829–39. Epub 20130108. doi: 10.1002/hep.26094. PubMed PMID: 23081697.

20. Boivin D, Lamy S, Lord-Dufour S, Jackson J, Beaulieu E, Côté M, et al. Antiproliferative and antioxidant activities of common vegetables: A comparative study. Food Chemistry. 2009;112(2):374–80. doi: 10.1016/j.foodchem.2008.05.084.

21. Corbett TH, Griswold DP, Jr., Roberts BJ, Peckham JC, Schabel FM, Jr. Tumor induction relationships in development of transplantable cancers of the colon in mice for chemotherapy assays, with a note on carcinogen structure. Cancer Res. 1975;35(9):2434–9. Epub 1975/09/11. PubMed PMID: 1149045.

22. Almeida JL, Hill CR, Cole KD. Mouse cell line authentication. Cytotechnology. 2014;66(1):133–47. Epub 20130222. doi: 10.1007/s10616-013-9545-7. PubMed PMID: 23430347; PubMed Central PMCID: PMCPMC3886540.

23. Sakaue-Sawano A, Kurokawa H, Morimura T, Hanyu A, Hama H, Osawa H, et al. Visualizing spatiotemporal dynamics of multicellular cell-cycle progression. Cell. 2008;132(3):487–98. doi: 10.1016/j.cell.2007.12.033. PubMed PMID: 18267078.

24. Bankhead P, Loughrey MB, Fernández JA, Dombrowski Y, McArt DG, Dunne PD, et al. QuPath: Open source software for digital pathology image analysis. Sci Rep. 2017;7(1):16878. Epub 20171204. doi: 10.1038/s41598-017-17204-5. PubMed PMID: 29203879; PubMed Central PMCID: PMCPMC5715110.

25. Team P. RStudio. PBC, Boston, MA; 2025.

26. Ehrchen JM, Roth J, Barczyk-Kahlert K. More Than Suppression: Glucocorticoid Action on Monocytes and Macrophages. Front Immunol. 2019;10:2028. Epub 20190827. doi: 10.3389/fimmu.2019.02028. PubMed PMID: 31507614; PubMed Central PMCID: PMCPMC6718555.

27. Ramakers C, Ruijter JM, Deprez RH, Moorman AF. Assumption-free analysis of quantitative real-time polymerase chain reaction (PCR) data. Neuroscience letters. 2003;339(1):62–6. Epub 2003/03/06. PubMed PMID: 12618301.

28. Pfaffl MW. A new mathematical model for relative quantification in real-time RT-PCR. Nucleic acids research. 2001;29(9):e45. Epub 2001/05/09. PubMed PMID: 11328886; PubMed Central PMCID: PMCPmc55695.

29. Godon JJ, Zumstein E, Dabert P, Habouzit F, Moletta R. Molecular microbial diversity of an anaerobic digestor as determined by small-subunit rDNA sequence analysis. Appl Environ Microbiol. 1997;63(7):2802–13. doi: 10.1128/aem.63.7.2802-2813.1997. PubMed PMID: 9212428; PubMed Central PMCID: PMCPMC168577.

30. Klindworth A, Pruesse E, Schweer T, Peplies J, Quast C, Horn M, et al. Evaluation of general 16S ribosomal RNA gene PCR primers for classical and next-generation sequencing-based diversity studies. Nucleic acids research. 2013;41(1):e1. Epub 20120828. doi: 10.1093/nar/gks808. PubMed PMID: 22933715; PubMed Central PMCID: PMCPMC3592464.

31. Berry D, Ben Mahfoudh K, Wagner M, Loy A. Barcoded primers used in multiplex amplicon pyrosequencing bias amplification. Appl Environ Microbiol. 2011;77(21):7846–9. Epub 20110902. doi: 10.1128/aem.05220-11. PubMed PMID: 21890669; PubMed Central PMCID: PMCPMC3209180.

32. Chen S, Zhou Y, Chen Y, Gu J. fastp: an ultra-fast all-in-one FASTQ preprocessor. Bioinformatics. 2018;34(17):i884–i90. doi: 10.1093/bioinformatics/bty560. PubMed PMID: 30423086; PubMed Central PMCID: PMCPMC6129281.

33. Bolyen E, Rideout JR, Dillon MR, Bokulich NA, Abnet CC, Al-Ghalith GA, et al. Reproducible, interactive, scalable and extensible microbiome data science using QIIME 2. Nat Biotechnol. 2019;37(8):852–7. doi: 10.1038/s41587-019-0209-9. PubMed PMID: 31341288; PubMed Central PMCID: PMCPMC7015180.

34. Callahan BJ, McMurdie PJ, Rosen MJ, Han AW, Johnson AJ, Holmes SP. DADA2: High-resolution sample inference from Illumina amplicon data. Nat Methods. 2016;13(7):581–3. Epub 20160523. doi: 10.1038/nmeth.3869. PubMed PMID: 27214047; PubMed Central PMCID: PMCPMC4927377.

35. Quast C, Pruesse E, Yilmaz P, Gerken J, Schweer T, Yarza P, et al. The SILVA ribosomal RNA gene database project: improved data processing and web-based tools. Nucleic acids research. 2013;41(Database issue):D590–6. Epub 20121128. doi: 10.1093/nar/gks1219. PubMed PMID: 23193283; PubMed Central PMCID: PMCPMC3531112.

36. Douglas GM, Maffei VJ, Zaneveld JR, Yurgel SN, Brown JR, Taylor CM, et al. PICRUSt2 for prediction of metagenome functions. Nat Biotechnol. 2020;38(6):685–8. doi: 10.1038/s41587-020-0548-6. PubMed PMID: 32483366; PubMed Central PMCID: PMCPMC7365738.

37. Temaj G, Saha S, Chichiarelli S, Telkoparan-Akillilar P, Nuhii N, Hadziselimovic R, et al. Polyploid giant cancer cells: Underlying mechanisms, signaling pathways, and therapeutic strategies. Critical Reviews in Oncology/Hematology. 2025;213:104802. doi: 10.1016/j.critrevonc.2025.104802.

38. Auguste P, Fallavollita L, Wang N, Burnier J, Bikfalvi A, Brodt P. The Host Inflammatory Response Promotes Liver Metastasis by Increasing Tumor Cell Arrest and Extravasation. The American Journal of Pathology. 2007;170(5):1781–92. doi: 10.2353/ajpath.2007.060886.

39. Cassetta L, Pollard JW. A timeline of tumour-associated macrophage biology. Nat Rev Cancer. 2023;23(4):238–57. Epub 20230215. doi: 10.1038/s41568-022-00547-1. PubMed PMID: 36792751.

40. Meyers M, Stoffels CBA, Frache G, Letellier E, Feucherolles M. Microbiome in cancer metastasis: biological insights and emerging spatial omics methods. Front Cell Infect Microbiol. 2025;15:1559870. Epub 20250604. doi: 10.3389/fcimb.2025.1559870. PubMed PMID: 40535543; PubMed Central PMCID: PMCPMC12174425.

41. Pleguezuelos-Manzano C, Puschhof J, Rosendahl Huber A, van Hoeck A, Wood HM, Nomburg J, et al. Mutational signature in colorectal cancer caused by genotoxic pks(+) E. coli. Nature. 2020;580(7802):269–73. Epub 20200227. doi: 10.1038/s41586-020-2080-8. PubMed PMID: 32106218; PubMed Central PMCID: PMCPMC8142898.

42. Liu QL, Zhou H, Wang Z, Chen Y. Exploring the role of gut microbiota in colorectal liver metastasis through the gut-liver axis. Front Cell Dev Biol. 2025;13:1563184. Epub 20250313. doi: 10.3389/fcell.2025.1563184. PubMed PMID: 40181829; PubMed Central PMCID: PMCPMC11965903.

43. Osswald A, Wortmann E, Wylensek D, Kuhls S, Coleman OI, Peuker K, et al. Secondary bile acid production by gut bacteria promotes Western diet-associated colorectal cancer. Gut. 2025. Epub 20251218. doi: 10.1136/gutjnl-2024-332243. PubMed PMID: 41412727.

44. Caserta S, Genovese C, Cicero N, Toscano V, Gangemi S, Allegra A. The Interplay between Medical Plants and Gut Microbiota in Cancer. Nutrients. 2023;15(15). Epub 20230726. doi: 10.3390/nu15153327. PubMed PMID: 37571264; PubMed Central PMCID: PMCPMC10421419.

45. Lagkouvardos I, Kläring K, Heinzmann SS, Platz S, Scholz B, Engel KH, et al. Gut metabolites and bacterial community networks during a pilot intervention study with flaxseeds in healthy adult men. Mol Nutr Food Res. 2015;59(8):1614–28. Epub 20150605. doi: 10.1002/mnfr.201500125. PubMed PMID: 25988339.

46. Ma J, Shi Q, Cui G, Sheng H, Botuyan MV, Zhou Y, et al. SPOP mutation induces replication over-firing by impairing Geminin ubiquitination and triggers replication catastrophe upon ATR inhibition. Nat Commun. 2021;12(1):5779. Epub 20211001. doi: 10.1038/s41467-021-26049-6. PubMed PMID: 34599168; PubMed Central PMCID: PMCPMC8486843.

47. Toledo L, Neelsen KJ, Lukas J. Replication Catastrophe: When a Checkpoint Fails because of Exhaustion. Molecular Cell. 2017;66(6):735–49. doi: 10.1016/j.molcel.2017.05.001.

48. Lyubitelev A, Studitsky V. Inhibition of Cancer Development by Natural Plant Polyphenols: Molecular Mechanisms. Int J Mol Sci. 2023;24(13). Epub 20230626. doi: 10.3390/ijms241310663. PubMed PMID: 37445850; PubMed Central PMCID: PMCPMC10341686.

49. Kapinova A, Kubatka P, Liskova A, Baranenko D, Kruzliak P, Matta M, et al. Controlling metastatic cancer: the role of phytochemicals in cell signaling. J Cancer Res Clin Oncol. 2019;145(5):1087–109. Epub 20190322. doi: 10.1007/s00432-019-02892-5. PubMed PMID: 30903319.

50. Li J, Yan Z, Li H, Shi Q, Ahire V, Zhang S, et al. The Phytochemical Scoulerine Inhibits Aurora Kinase Activity to Induce Mitotic and Cytokinetic Defects. J Nat Prod. 2021;84(8):2312–20. Epub 20210818. doi: 10.1021/acs.jnatprod.1c00429. PubMed PMID: 34406008.

51. Salmela AL, Pouwels J, Varis A, Kukkonen AM, Toivonen P, Halonen PK, et al. Dietary flavonoid fisetin induces a forced exit from mitosis by targeting the mitotic spindle checkpoint. Carcinogenesis. 2009;30(6):1032–40. Epub 20090424. doi: 10.1093/carcin/bgp101. PubMed PMID: 19395653; PubMed Central PMCID: PMCPMC2691139.

52. Stivers NS, Islam A, Reyes-Reyes EM, Casson LK, Aponte JC, Vaisberg AJ, et al. Plagiochiline A Inhibits Cytokinetic Abscission and Induces Cell Death. Molecules. 2018;23(6). Epub 20180612. doi: 10.3390/molecules23061418. PubMed PMID: 29895732; PubMed Central PMCID: PMCPMC6099941.

53. López-Gómez L, Uranga JA. Polyphenols in the Prevention and Treatment of Colorectal Cancer: A Systematic Review of Clinical Evidence. Nutrients. 2024;16(16). Epub 20240816. doi: 10.3390/nu16162735. PubMed PMID: 39203871; PubMed Central PMCID: PMCPMC11357634.

54. Okpoghono J, Isoje EF, Igbuku UA, Ekayoda O, Omoike GO, Adonor TO, et al. Natural polyphenols: A protective approach to reduce colorectal cancer. Heliyon. 2024;10(11):e32390. doi: 10.1016/j.heliyon.2024.e32390.

55. Narayanan NK, Kunimasa K, Yamori Y, Mori M, Mori H, Nakamura K, et al. Antitumor activity of melinjo (Gnetum gnemon L.) seed extract in human and murine tumor models in vitro and in a colon-26 tumor-bearing mouse model in vivo. Cancer Med. 2015;4(11):1767–80. Epub 20150926. doi: 10.1002/cam4.520. PubMed PMID: 26408414; PubMed Central PMCID: PMCPMC4674003.

56. Yuan J-H, Li Y-Q, Yang X-Y. Inhibition of Epigallocatechin Gallate on Orthotopic Colon Cancer by Upregulating the Nrf2-UGT1A Signal Pathway in Nude Mice. Pharmacology. 2007;80(4):269–78. doi: 10.1159/000106447.

57. Wang H, Wu X, Lezmi S, Li Q, Helferich WG, Xu Y, et al. Extract of Ginkgo biloba exacerbates liver metastasis in a mouse colon cancer Xenograft model. BMC Complementary and Alternative Medicine. 2017;17(1):516. doi: 10.1186/s12906-017-2014-7.

58. Crusz SM, Balkwill FR. Inflammation and cancer: advances and new agents. Nat Rev Clin Oncol. 2015;12(10):584–96. Epub 20150630. doi: 10.1038/nrclinonc.2015.105. PubMed PMID: 26122183.

59. Shakoor H, Feehan J, Apostolopoulos V, Platat C, Al Dhaheri AS, Ali HI, et al. Immunomodulatory Effects of Dietary Polyphenols. Nutrients. 2021;13(3). Epub 20210225. doi: 10.3390/nu13030728. PubMed PMID: 33668814; PubMed Central PMCID: PMCPMC7996139.

60. Pyonteck SM, Akkari L, Schuhmacher AJ, Bowman RL, Sevenich L, Quail DF, et al. CSF-1R inhibition alters macrophage polarization and blocks glioma progression. Nat Med. 2013;19(10):1264–72. Epub 20130922. doi: 10.1038/nm.3337. PubMed PMID: 24056773; PubMed Central PMCID: PMCPMC3840724.

61. Qian BZ, Li J, Zhang H, Kitamura T, Zhang J, Campion LR, et al. CCL2 recruits inflammatory monocytes to facilitate breast-tumour metastasis. Nature. 2011;475(7355):222–5. Epub 20110608. doi: 10.1038/nature10138. PubMed PMID: 21654748; PubMed Central PMCID: PMCPMC3208506.

62. Schmid MC, Avraamides CJ, Foubert P, Shaked Y, Kang SW, Kerbel RS, et al. Combined blockade of integrin-α4β1 plus cytokines SDF-1α or IL-1β potently inhibits tumor inflammation and growth. Cancer Res. 2011;71(22):6965–75. Epub 20110923. doi: 10.1158/0008-5472.Can-11-0588. PubMed PMID: 21948958; PubMed Central PMCID: PMCPMC3249446.

63. Morgan RA. Human tumor xenografts: the good, the bad, and the ugly. Mol Ther. 2012;20(5):882–4. doi: 10.1038/mt.2012.73. PubMed PMID: 22549804; PubMed Central PMCID: PMCPMC3345993.

## Supplementary References

1. Chen, S., Zhou, Y., Chen, Y., & Gu, J. (2018). fastp: an ultra-fast all-in-one FASTQ preprocessor. Bioinformatics, 34(17), i884–i890.

2. Bolyen, E., Rideout, J. R., Dillon, M. R., Bokulich, N. A., Abnet, C. C., Al-Ghalith, G. A., Caporaso, J. G. (2019). Reproducible, interactive, scalable and extensible microbiome data science using QIIME 2. Nature biotechnology, 37(8), 852–857.

3. Martin, M. (2011). Cutadapt removes adapter sequences from high-throughput sequencing reads. EMBnet. journal, 17(1), 10–12.

4. Callahan, B. J., McMurdie, P. J., Rosen, M. J., Han, A. W., Johnson, A. J. A., & Holmes, S. P. (2016). DADA2: High-resolution sample inference from Illumina amplicon data. Nature methods, 13(7), 581–583.

5. Quast, C., Pruesse, E., Yilmaz, P., Gerken, J., Schweer, T., Yarza, P., … & Glöckner, F. O. (2012). The SILVA ribosomal RNA gene database project: improved data processing and web-based tools. Nucleic acids research, 41(D1), D590–D596.

6. Robeson, M. S., O’Rourke, D. R., Kaehler, B. D., Ziemski, M., Dillon, M. R., Foster, J. T., & Bokulich, N. A. (2021). RESCRIPt: Reproducible sequence taxonomy reference database management. PLoS computational biology, 17(11), e1009581.

7. McMurdie, P. J., & Holmes, S. (2013). phyloseq: an R package for reproducible interactive analysis and graphics of microbiome census data. PloS one, 8(4), e61217.

8. Oksanen J, Simpson G, Blanchet F, Kindt R, Legendre P, Minchin P, O’Hara R, Solymos P, Stevens M, Szoecs E, Wagner H, Barbour M, Bedward M, Bolker B, Borcard D, Borman T, Carvalho G, Chirico M, De Caceres M, Durand S, Evangelista H, FitzJohn R, Friendly M, Furneaux B, Hannigan G, Hill M, Lahti L, Martino C, McGlinn D, Ouellette M, Ribeiro Cunha E, Smith T, Stier A, Ter Braak C, Weedon J (2025). _vegan: Community Ecology Package_. doi:10.32614/CRAN.package.vegan <10.32614/CRAN.package.vegan>, R package version 2.7-1, <https://CRAN.R-project.org/package=vegan>.

9. Martinez Arbizu P (2017). _pairwiseAdonis: Pairwise Multilevel Comparison using Adonis_. R package version 0.4.1, commit cb190f7668a0c82c0b0853927db239e7b9ec3e83, <https://github.com/pmartinezarbizu/pairwiseAdonis>.

10. Reitmeier, S., Hitch, T.C., Treichel, N., Fikas, N., Hausmann, B., Ramer-Tait, A.E., Neuhaus, K., Berry, D., Haller, D., Lagkouvardos, I. and Clavel, T. (2021). Handling of spurious sequences affects the outcome of high-throughput 16S rRNA gene amplicon profiling. ISME communications, 1(1), p.31.

11. Gu, Z., Eils, R., & Schlesner, M. (2016). Complex heatmaps reveal patterns and correlations in multidimensional genomic data. Bioinformatics, 32(18), 2847–2849.

