## Supplemental Figures and Tables for "A mixture of plant polyphenols unexpectedly aggravates liver metastasis of colorectal cancer in mice"

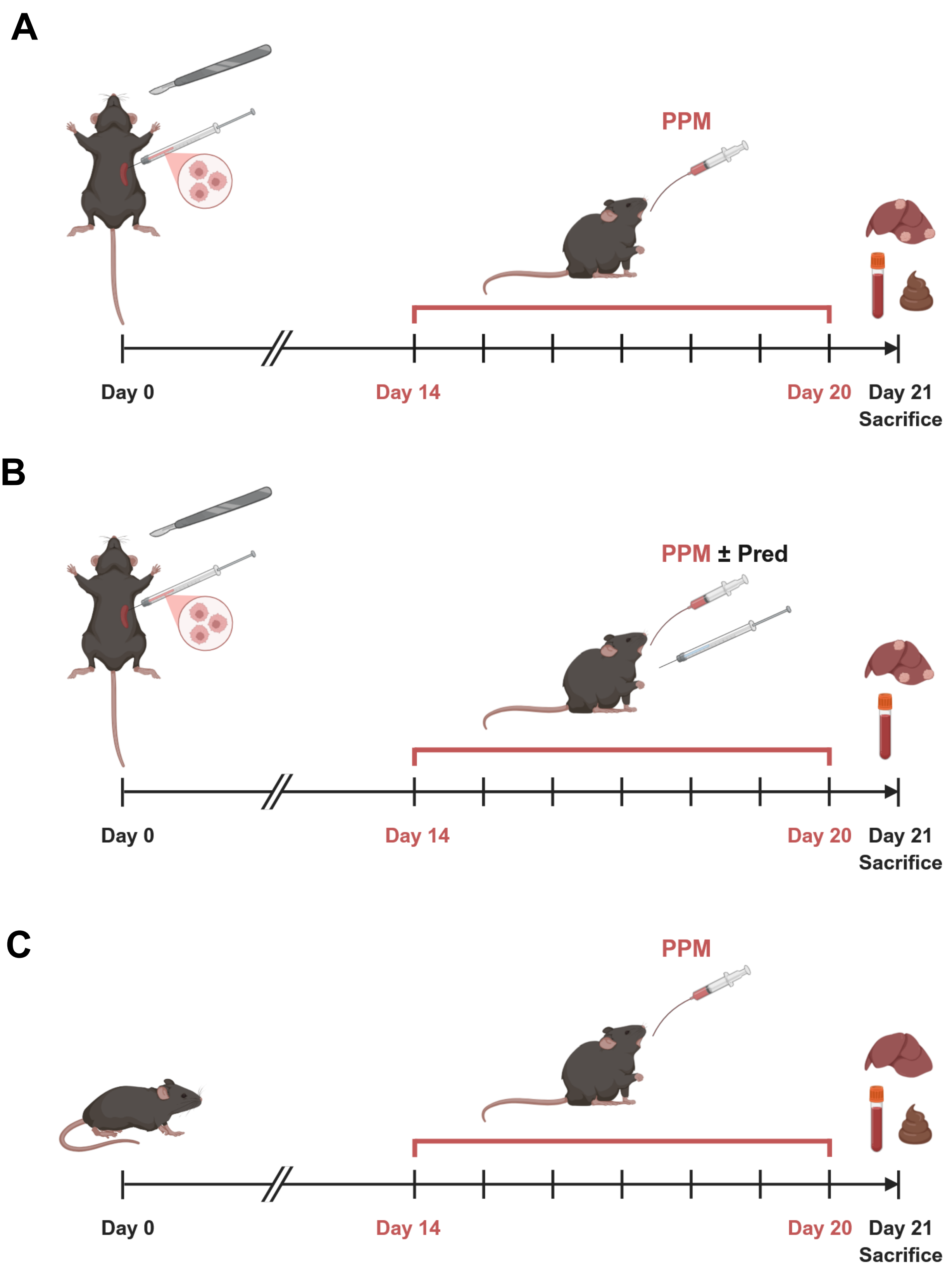
A
PPM
Day 0
Day 14
Day 20
Day 21
Sacrifice
B
PPM ± Pred
Day 0
Day 14
Day 20
Day 21
Sacrifice
C
PPM
Day 0
Day 14
Day 20
Day 21
Sacrifice

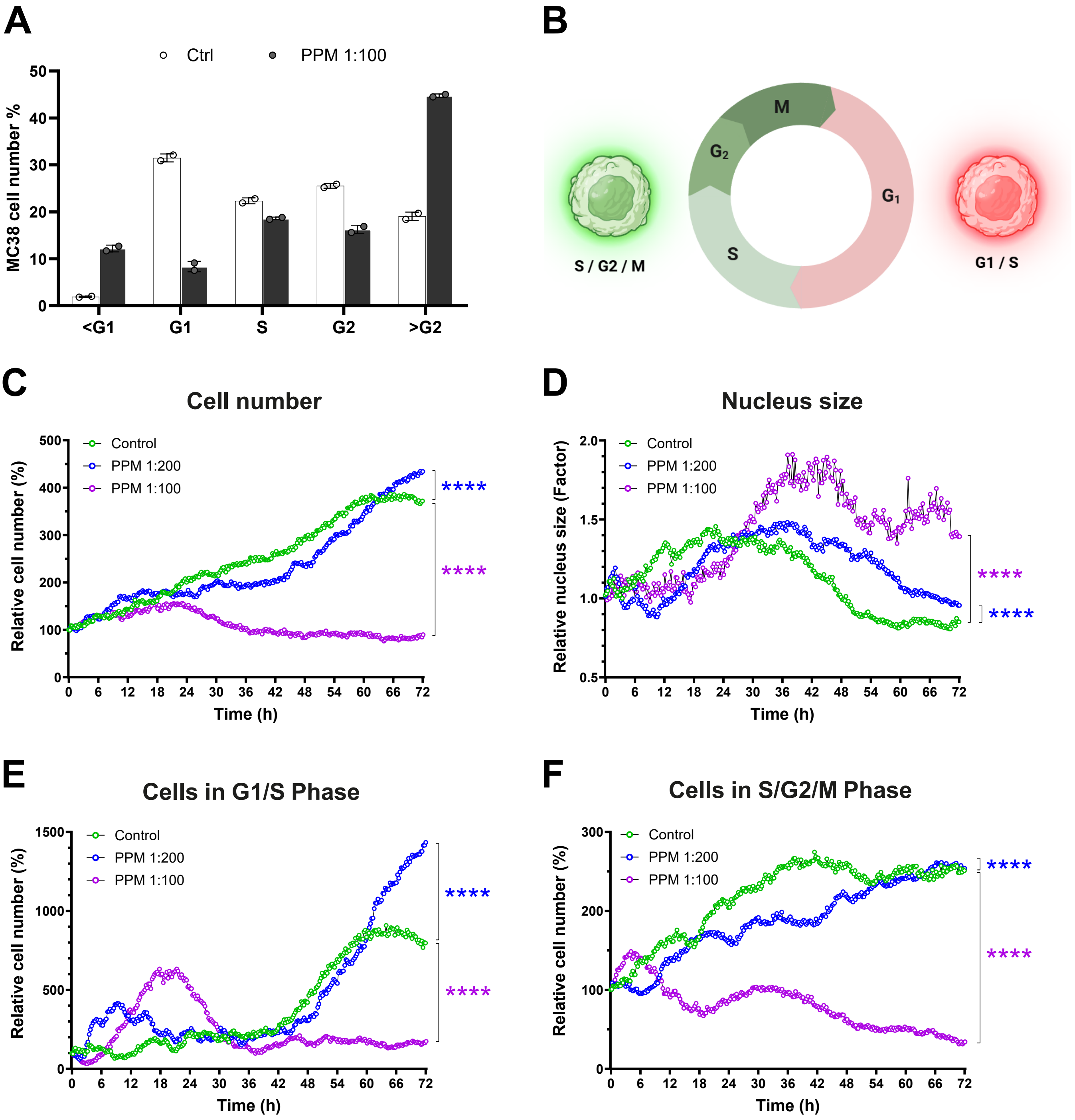


G

| 48 hours | | |
|---|---|---|
| **Comparison** | **Significance** | ***p* value** |
| Control / PPM 1:200 | ** | 0.0085 |
| Control / PPM 1:100 | *** | 0.0004 |

| 72 hours | | |
|---|---|---|
| **Comparison** | **Significance** | ***p* value** |
| Control / PPM 1:200 | *** | 0.0002 |
| Control / PPM 1:100 | **** | <0.0001 |

| 96 hours | | |
|---|---|---|
| **Comparison** | **Significance** | ***p* value** |
| Control / PPM 1:200 | **** | <0.0001 |
| Control / PPM 1:100 | **** | <0.0001 |

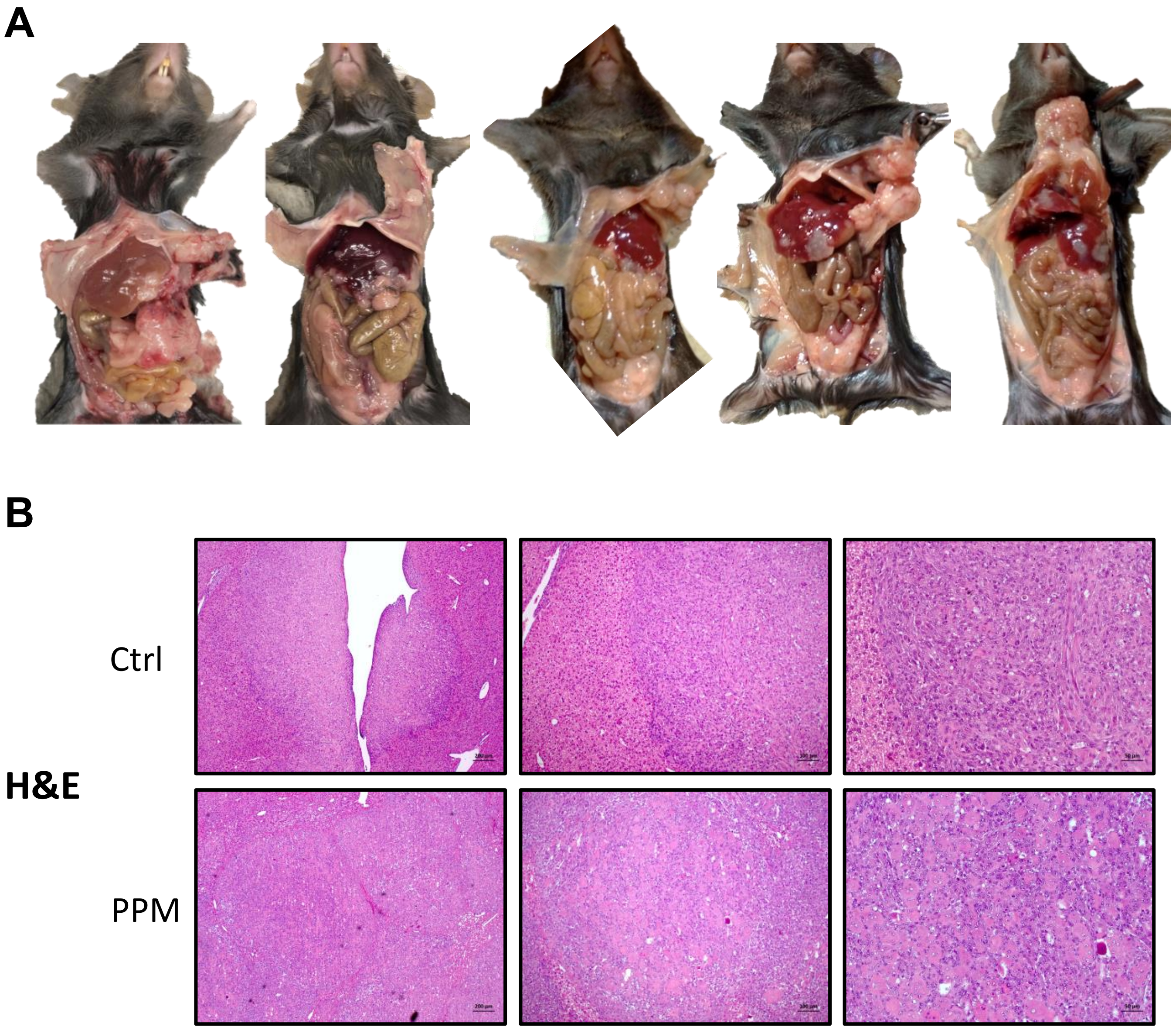
A
B
Ctrl
H&E
PPM

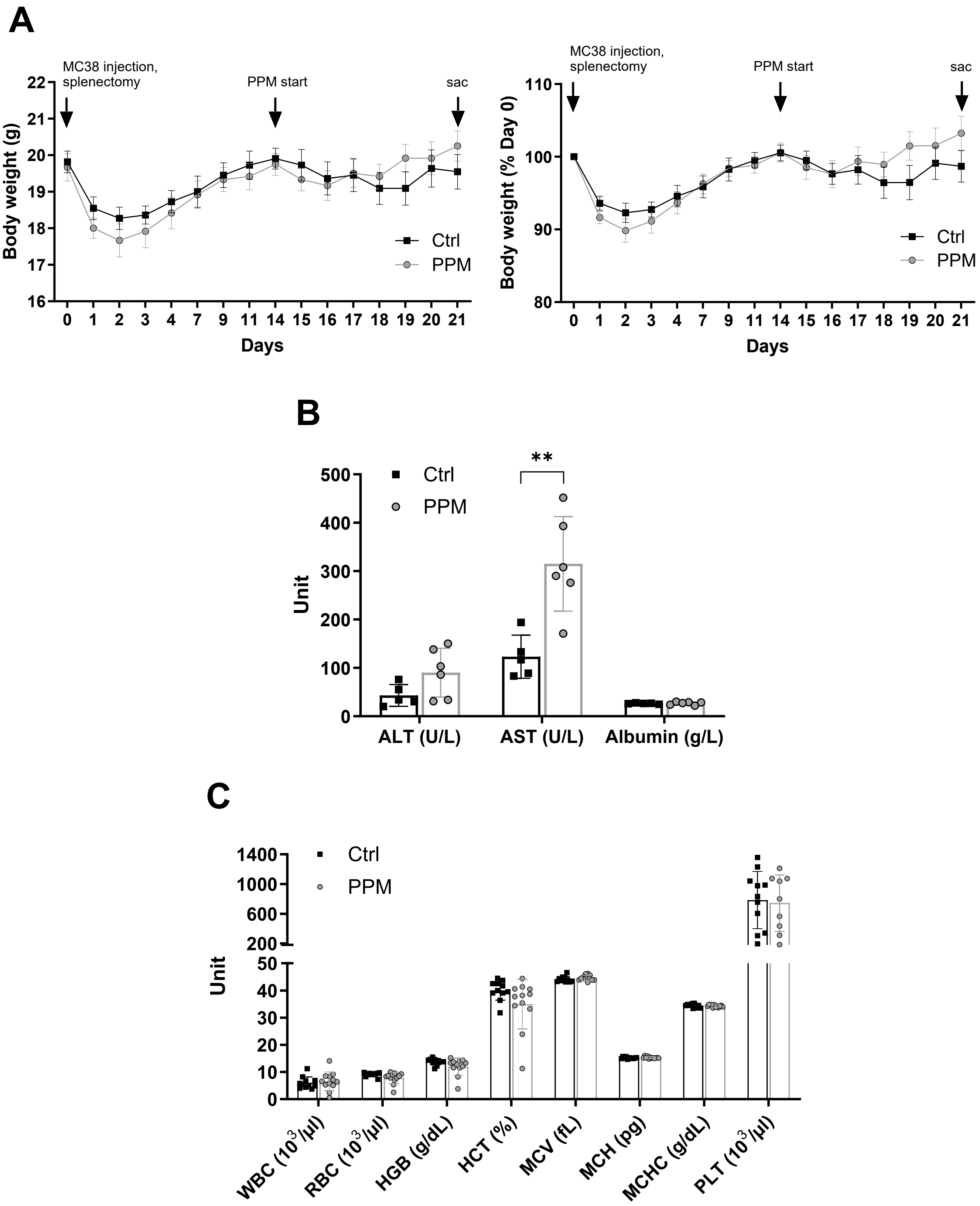
A
MC38 injection, splenectomy
PPM start
sac
Body weight (g)
Days
Ctrl
PPM
Body weight (% Day 0)
B
Unit
**
ALT (U/L)
AST (U/L)
Albumin (g/L)
C
WBC ($10^3$/µl)
RBC ($10^3$/µl)
HGB (g/dL)
HCT (%)
MCV (fL)
MCH (pg)
MCHC (g/dL)
PLT ($10^3$/µl)

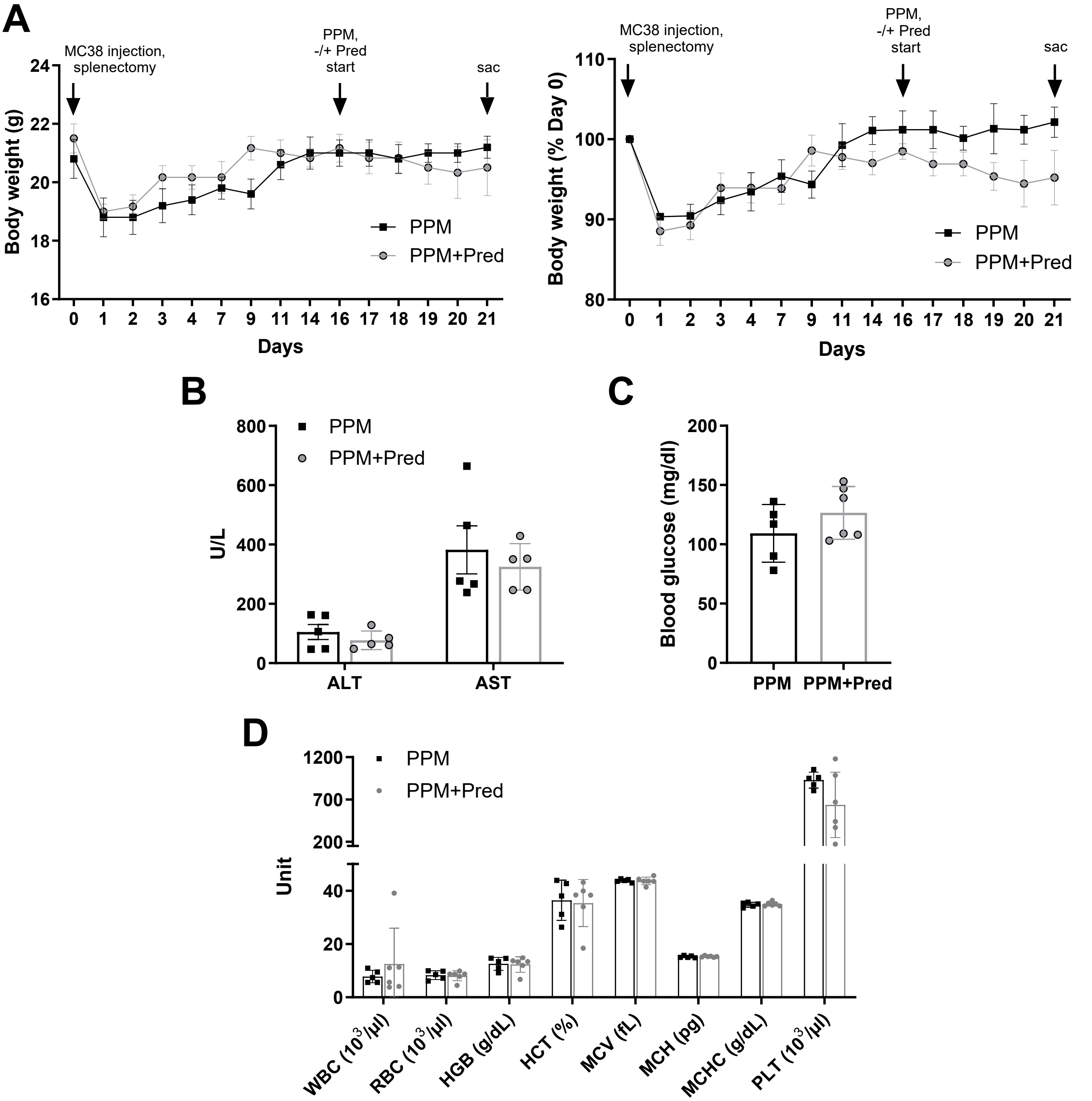
A
MC38 injection, splenectomy
PPM, -/+ Pred start
sac
Body weight (g)
Days
PPM
PPM+Pred
Body weight (% Day 0)
B
U/L
ALT
AST
C
Blood glucose (mg/dl)
D
Unit
WBC (10^3/μl)
RBC (10^3/μl)
HGB (g/dL)
HCT (%)
MCV (fL)
MCH (pg)
MCHC (g/dL)
PLT (10^3/μl)

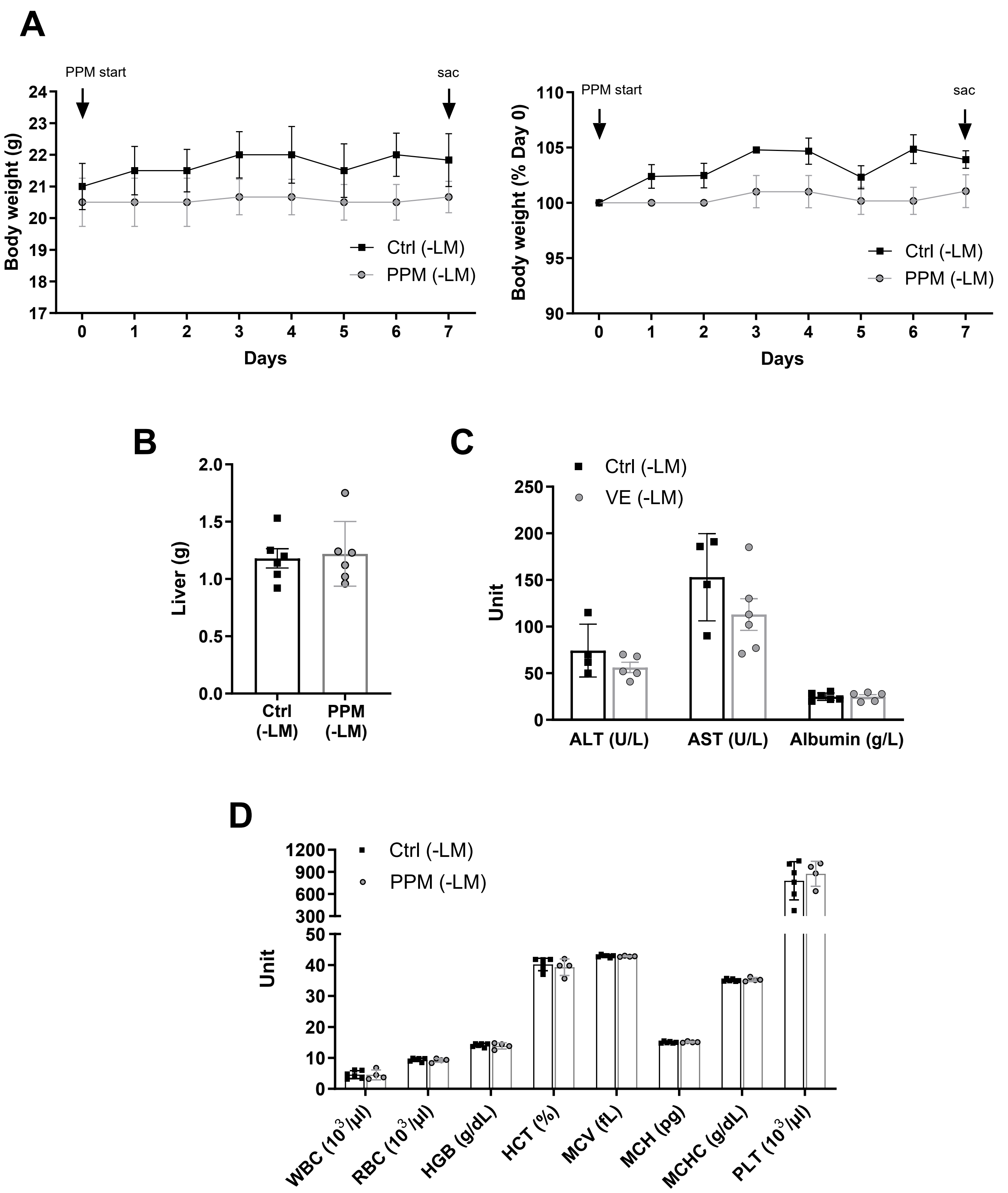
A
PPM start
sac
Body weight (g)
Days
Ctrl (-LM)
PPM (-LM)
Body weight (% Day 0)
B
Liver (g)
Ctrl (-LM)
PPM (-LM)
C
Ctrl (-LM)
VE (-LM)
Unit
ALT (U/L)
AST (U/L)
Albumin (g/L)
D
Ctrl (-LM)
PPM (-LM)
Unit
WBC (10³/µl)
RBC (10³/µl)
HGB (g/dL)
HCT (%)
MCV (fL)
MCH (pg)
MCHC (g/dL)
PLT (10³/µl)

A

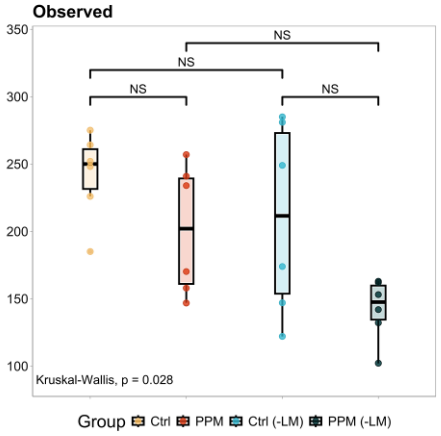


B

**Bray-Curtis pairwise**

| Comparison | F | Pr(>F) |
|---|---|---|
| PPM vs Ctrl | 1.419133 | 0.073 |
| PPM vs Ctrl (-LM) | 1.509729 | 0.126 |
| PPM vs PPM (-LM) | 3.076307 | 0.008 |
| Ctrl vs Ctrl (-LM) | 2.752822 | 0.001 |
| Ctrl vs PPM (-LM) | 6.571154 | 0.001 |
| Ctrl (-LM) vs PPM (-LM) | 1.734046 | 0.115 |

C

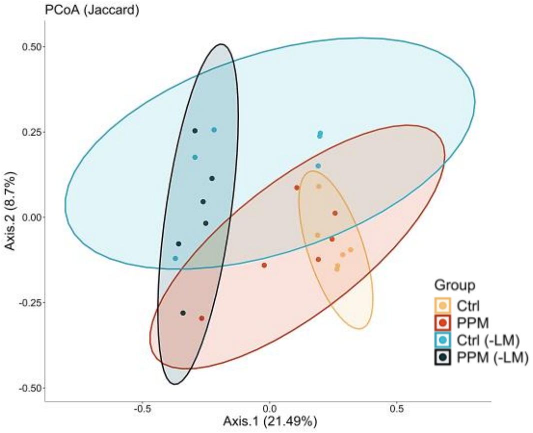


**Jaccard pairwise**

| Comparison | F | Pr(>F) |
|---|---|---|
| PPM vs Ctrl | 1.215826 | 0.130 |
| PPM vs Ctrl (-LM) | 1.362220 | 0.120 |
| PPM vs PPM (-LM) | 2.290017 | 0.010 |
| Ctrl vs Ctrl (-LM) | 2.083176 | 0.002 |
| Ctrl vs PPM (-LM) | 4.016169 | 0.002 |
| Ctrl (-LM) vs PPM (-LM) | 1.406035 | 0.131 |
